# Robustness of ancestral sequence reconstruction to among-site evolutionary heterogeneity and epistasis

**DOI:** 10.1101/2024.12.20.629812

**Authors:** Ricardo Muñiz-Trejo, Yeonwoo Park, Joseph W. Thornton

## Abstract

Ancestral sequence reconstruction (ASR) is typically performed using homogeneous evolutionary models, which assume that the same substitution propensities affect all sites and lineages. These assumptions are routinely violated: heterogeneous structural and functional constraints favor different amino acid states at different sites, and these constraints often change among lineages as epistatic substitutions accrue at other sites. To evaluate how realistic violations of the homogeneity assumption affect ASR, we developed site-specific substitution models and parameterized them using data from deep mutational scanning experiments on three protein families; we then used these models to perform ASR on the empirical alignments and on alignments simulated under heterogeneous conditions derived from the experiments. Extensive among-site and -lineage heterogeneity is present in these datasets, but the sequences reconstructed from empirical alignments are almost identical, irrespective of whether heterogeneous or homogeneous models are used for ASR. The rare differences occur primarily when phylogenetic signal is weak – at fast-evolving sites and nodes connected by long branches. When ASR is performed on simulated data, errors in the reconstructed sequences become more likely as branch lengths increase, but incorporating heterogeneity into the model does not improve accuracy. These data establish that ASR is robust to unincorporated realistic forms of evolutionary heterogeneity, because the primary determinant of ASR is phylogenetic signal, not the substitution model. The best way to improve accuracy is therefore not to develop more elaborate models but to apply ASR to densely sampled alignments that maximize phylogenetic signal at the nodes of interest.

## INTRODUCTION

Ancestral sequence reconstruction (ASR) has become an important strategy in molecular evolution for experimentally testing hypotheses about the properties of ancient proteins and the genetic and biochemical mechanisms by which those properties evolved during history (Thornton 2004; Liberles 2007; Merkl and Sterner 2016; Hochberg and Thornton 2017; Starr et al. 2020; Mascotti 2022). ASR finds the most probable ancestral sequence at any node on a phylogenetic tree, given an alignment of extant proteins, the phylogeny, and a stochastic substitution model that describes the relative rates of sequence change among the possible amino acid or nucleotide states (Yang et al. 1995). Previous research has evaluated the robustness of inferred ancestral sequences to using different alignment methods (Vialle et al. 2018) and to uncertainty about the phylogenetic tree (Hanson-Smith et al. 2010; Groussin et al. 2015). Here we address the sensitivity of ASR to model misspecification – the inevitable mismatch between the substitution model used for ASR and the true processes of historical sequence evolution. We examine two forms of evolutionary complexity that are seldom incorporated into substitution models for ASR: differences among sites in the equilibrium frequencies of amino acids and among-lineage differences in these frequencies.

The vast majority of ancestral reconstructions have been performed using site-homogeneous models, in which all amino acid sites in the protein have the same vector of expected frequencies of the 20 amino acids and the same matrix of relative rates of substitution between them. These models are typically estimated from very large sequence databases, with free parameters that represent an “average” set of frequencies and rates across all sites in a wide variety of proteins with different structures, functions, and histories (e.g., Jones et al. 1992; Whelan and Goldman 2001; Le and Gascuel 2008); equilibrium frequencies may also be estimated as the vector that best fits all sites in an alignment. The assumption of site-homogeneity is routinely violated (Naser-Khdour et al. 2019; Zou and Zhang 2019), because sites within a protein are subject to different structural and functional constraints (Kimura and Ohta 1974; Worth et al. 2009; Grahnen et al. 2011; Yeh et al. 2014). For example, hydrophobic amino acids are favored in the protein core but not at exposed surface sites, and the various amino acids have different propensities to be found in helices, sheets and loops because they favor different protein backbone angles. As a consequence, the frequencies of amino acid states and the rates at which each state is exchanged for the others are highly heterogeneous across sites (Halpern and Bruno 1998; Ashenberg et al. 2013; Echave et al. 2016). When tree topologies are inferred using homogeneous models that do not incorporate this among-site compositional heterogeneity, long-branch attraction artifacts can result (Lartillot et al. 2007; Feuda et al. 2017; Schrempf et al. 2020; Szánthó et al. 2023), and heterogeneous models that include compositional heterogeneity typically fit sequence data better and can reduce topological errors (Lartillot and Philippe 2004; Si Quang et al. 2008).

A second form of model violation is among-lineage compositional heterogeneity. If the effects of mutations depend on the genetic background into which they are introduced because of epistatic interactions, then the constraints that affect each site may change across the tree as sequences diverge at other sites. Epistatic interactions are clearly widespread during protein evolution (Breen et al. 2012; Gong et al. 2013; Bank et al. 2016; Park et al. 2022). Like site-specific compositional heterogeneity, unincorporated lineage-specific heterogeneity can also cause incorrect phylogenetic inferences (Foster 2004; Jayaswal et al. 2014).

The effect of these forms of model violation on ASR has not been thoroughly evaluated. Several studies have addressed how the choice among various site-homogeneous models affects the reconstructed sequences, observing only very small impacts on ancestral sequences and their accuracy (Pupko et al. 2007; Del Amparo and Arenas 2022; Sennett and Theobald 2023). These models are generally very similar to each other, with differences primarily attributable to averaging over different large protein datasets. The only work to directly address unincorporated compositional heterogeneity per se has been a series of computational studies in which sequence evolution was simulated under a simple biophysically-based model of site-heterogeneous effects on protein stability; the model used for ASR had very small effects on the inferred ancestral sequences or their predicted stabilities (Arenas et al. 2015; Arenas et al. 2017; Arenas and Bastolla 2020). We are aware of no studies that evaluated the effects of lineage-specific heterogeneity on ASR. It is therefore unknown how empirical forms of among-site and among-lineage compositional heterogeneity – which can reflect constraints on any aspect of protein biochemistry or function – affect the robustness and accuracy of sequences inferred by ancestral reconstruction.

Here we evaluate the effect of these forms of model violation on ASR. We incorporate realistic forms of compositional heterogeneity by using data from experimental deep mutational scanning (DMS) experiments, which directly characterize the functional effect of introducing every possible amino acid state at each site in a protein. For three example protein families, we used DMS data to parameterize site-specific heterogeneous substitution models and then used these models for ancestral sequence reconstruction. To assess robustness of ASR to including/excluding compositional heterogeneity, we compared ancestral sequences reconstructed using the site-specific DMS model versus those reconstructed using conventional site-homogeneous models. To assess the effect on ASR accuracy, we simulated sequence evolution of these protein families using the corresponding site-specific DMS model, and then compared the true ancestral sequences to those inferred using site-homogeneous and site-heterogeneous models. We also assessed the effect of among-lineage compositional heterogeneity by comparing evaluating accuracy and robustness of ASR using site-specific models parameterized using DMS experiments performed on distantly related proteins within the same family.

## RESULTS

### Experimentally informed site-specific substitution models

To incorporate realistic forms of among-site compositional heterogeneity into ancestral sequence reconstruction, we developed a site-specific substitution model that can be parameterized using data from DMS experiments that measure the effect of every possible amino acid mutation at every site in a protein. Conventional site-homogeneous models specify a 20 × 20 matrix of instantaneous substitution rates by which each amino acid *y* replaces an amino acid *x*, and this matrix applies to all sites in the protein. Our site-specific models specify a unique rate matrix for every site in the protein. Our approach follows that of Bloom (2014a, 2014b), who developed a site-specific model of nucleotide substitution which was parameterized using experimental data on the fitness effect of mutations at each site. The major modifications in our approach are to accommodate amino acid alignments (rather than nucleotide or codon alignments) and to incorporate DMS measurements of function rather than direct measurements of fitness.

Like Bloom’s, our model uses the formalism of Halpern and Bruno (1998), in which the rate of substitution from amino acid *x* to *y* is simply the rate of *x* mutating into *y*, multiplied by the probability of this mutation being fixed in the population (Fig. 1). Our model assumes that amino acid mutation rates are homogeneous across sites and are determined by nucleotide mutation rates acting through the standard genetic code; the nucleotide mutation rates are represented using a general time-reversible model. The probabilities of fixation are site-specific and are derived from the functional effects of each amino acid mutation measured in the DMS experiment. A simple sigmoid relationship is used to convert effects on function to fitness effects; this model represents purifying selection to maintain function, with fitness that plateaus at an upper bound (the wild-type fitness) as function increases, and plateaus at a lower bound of zero as function decreases (Chou et al. 2011; Bank 2022). Fitness effects determine fixation probabilities via the classic Kimura equation. Finally, we capture any unaccounted-for variation in the total rate of substitution among sites using a discrete gamma model (Yang 1994). The free parameters of the model – eight rate parameters of the nucleotide mutation model, three free parameters that determine the exact shape of the sigmoid curve, and the shape parameter of the gamma distribution – are estimated from the alignment data by maximum likelihood. This model was used for branch length optimization and ASR on a specified topology, not for identifying the topology itself.

**Figure 1.**
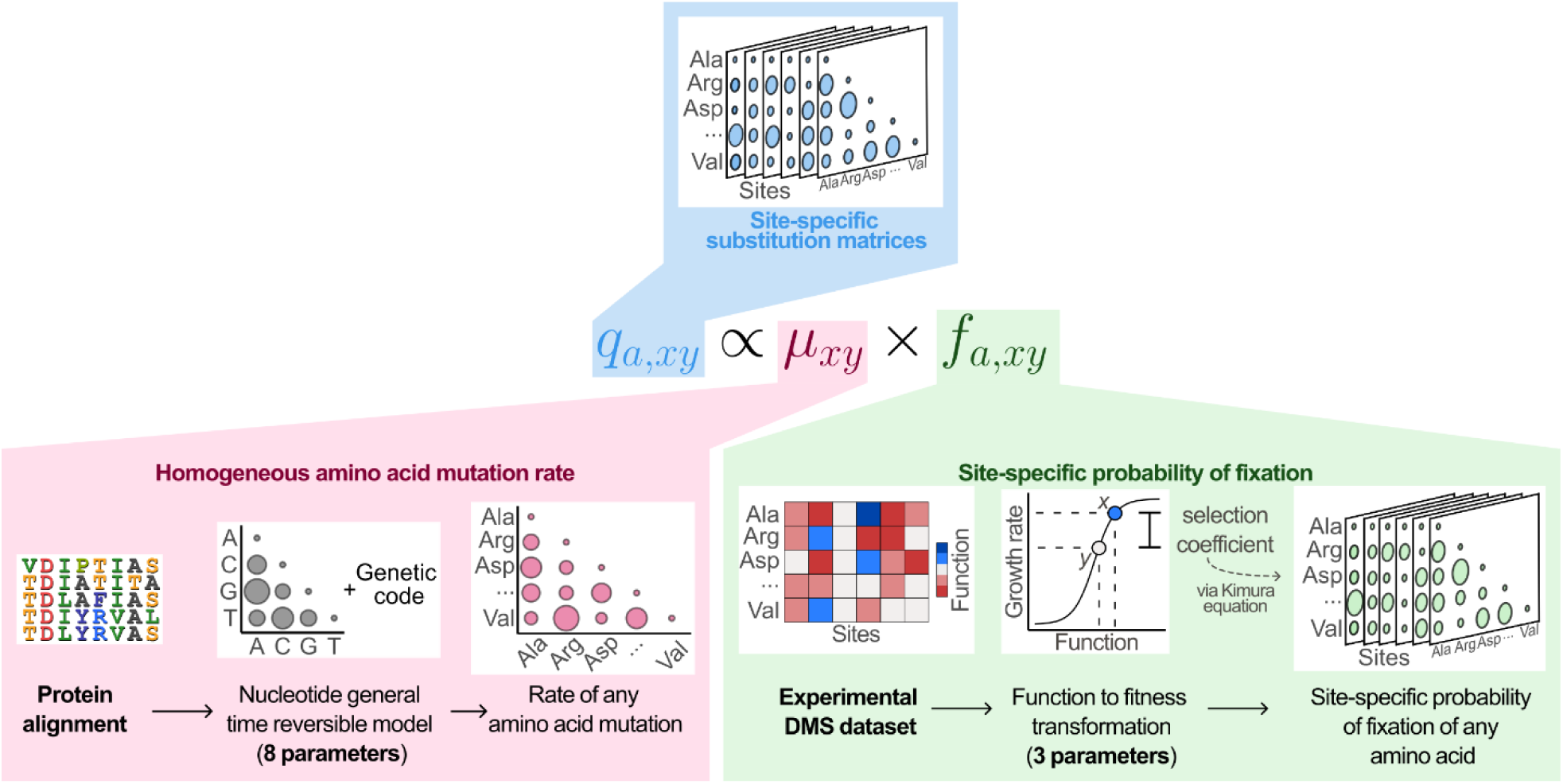
Graphical summary of site-specific substitution model generation. The site-specific substitution model specifies the instantaneous rate of substitution of any amino acid *x* by any other *y* at each site *a* (blue box). Each substitution rate is the product of the rate at which mutation produces an amino acid change (pink box) and the probability that the change will be fixed (green box). The mutation rate, which is homogeneous across sites, is parameterized from the sequence alignment assuming a general-time reversible model of nucleotide mutation and the standard genetic code. The fixation probability is parameterized by transforming experimental data on the functional effects of each amino acid state at each site using a simple logistic function-to-fitness relationship.

### Protein Systems Analyzed

We used our site-specific model to investigate the accuracy and robustness of ASR using three example protein families. These families were selected because they all have high-quality DMS datasets, well-resolved phylogenetic trees, and robust multiple sequence alignments. They differ dramatically in their biological functions, the types of phenotype measured, the rates at which they evolve, and the timescale encompassed by their phylogenies (Fig. 2A).

**Figure 2.**
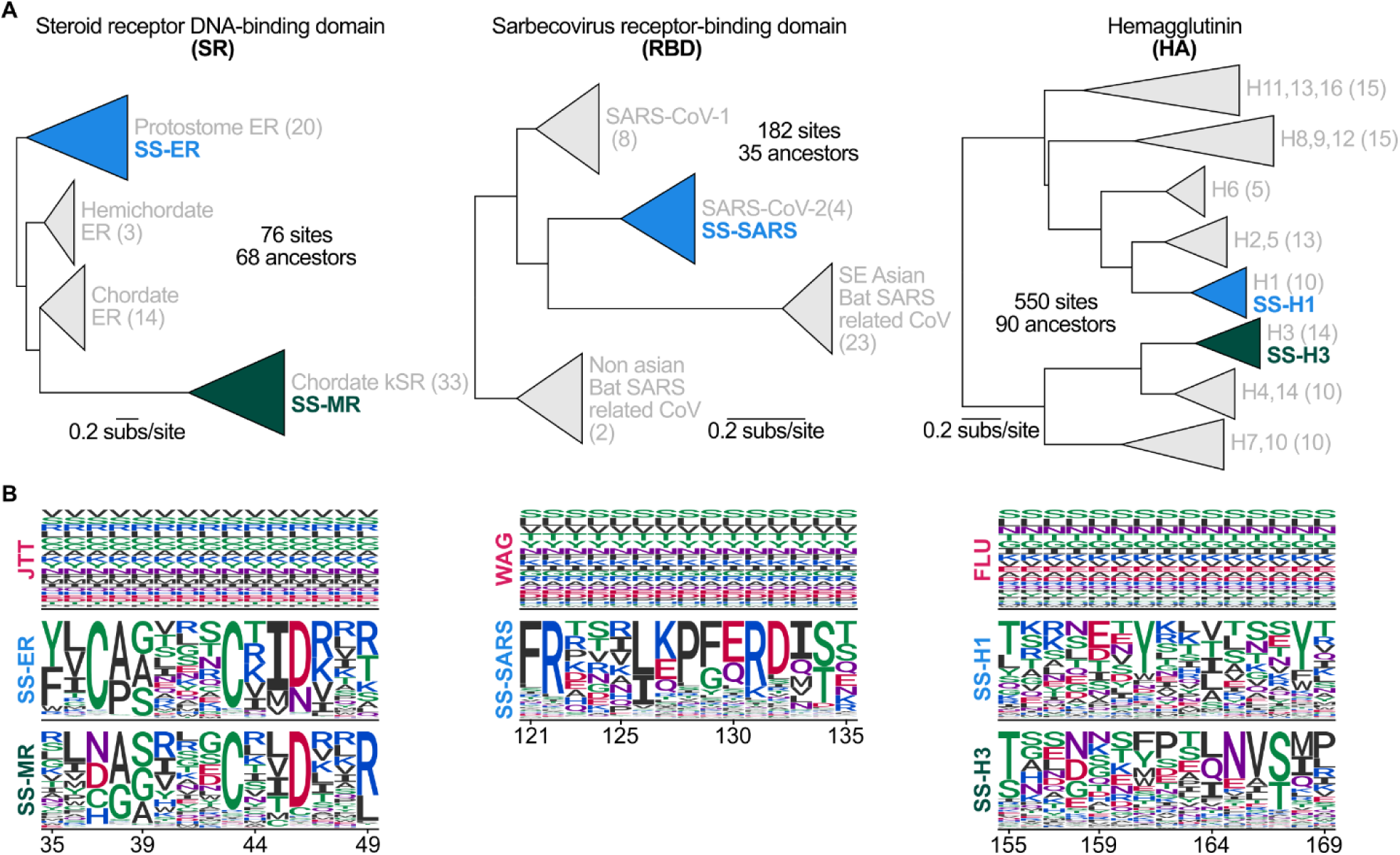
Protein families selected for analysis in this work. **(A)** Reduced phylogenetic trees of SR (left), RBD (middle), and HA (right). The proteins used for DMS that yield the best fit to the alignment are blue; the proteins used for DMS with the second-best fit, which was used for analyzing among-lineage compositional heterogeneity, are green. Number of terminal branches per clade in parenthesis. See Supplementary Figure S1-S3 for detailed topologies. **(C)** Examples of the differences in equilibrium frequencies estimated by the best-fitting site-homogeneous models (top row) versus the best-fitting site-specific models (bottom row). Only a subset of sites is shown, see Supp. Figs. S4-S6 for a more complete display of the site-specific empirical frequencies per model.

The first system is the DNA binding domain of steroid hormone receptors (SRs), a family of transcription factors that regulate many developmental and physiological processes in bilaterian animals (Beato et al. 1995; Mangelsdorf et al. 1995; Klinge 2018). Park et al. (2022) inferred the phylogeny of this protein family (Supp. Fig. S1) – the common ancestor of which existed >600 mya – and experimentally measured the effects of all amino acid replacements at each of the 76 sites in the DNA binding domain on DNA-binding using a bulk GFP reporter assay in yeast. The DMS experiment was performed using several extant and reconstructed ancestral proteins as backgrounds in which the mutant libraries were constructed. Here we fit site-specific models using the libraries constructed in two distantly related paralogs – estrogen receptor of the annelid *Capitella teleta* and the mineralocorticoid receptor of *Homo sapiens.* We refer to these fitted models as SS-ER and SS-MR, respectively.

The second system is the receptor-binding domain (RBD) of the Spike glycoprotein of Sarbecoviruses, which mediates binding of Spike to the ACE2 receptor and subsequent viral entry into mammalian cells (Walls et al. 2020). Starr et al. (2020) inferred the RBD phylogeny (Supp. Fig. S2) from SARS-CoV-1, SARS-CoV-2 and related viruses and used a yeast display system to assay the binding affinities to human ACE2 of variants of SARS-CoV-2 containing all amino acid replacements at all 182 sites of the RBD. We used these data to fit the site-specific substitution model for this protein (SS-SARS). This phylogeny spans a few decades of viral evolution but involves substantial sequence divergence because the protein evolves rapidly.

The third system, the influenza hemagglutinin (HA) protein, binds host glycoproteins and mediates membrane fusion into endosomes (Russell 2008; Gamblin et al. 2021). A multiple sequence alignment and phylogenetic tree were inferred for 90 HA family members representing 15 HA subtypes (Hilton & Bloom, 2018) (Supp. Fig. S3). Function was assessed by measuring the fitness effects of a library of all single-amino acid replacements at all 550 sites in the protein using a DMS assay that quantifies variant frequency pre- and post-viral passage. DMS libraries were engineered and assessed in two different backgrounds – the HA subtypes H1 (Doud and Bloom 2016) and H3 (Lee et al. 2018). We used both datasets to fit site-specific models (SS-H1 and SS-H3). Like the SARS RBD, the HA phylogeny spans several decades of rapid evolution.

For each system, we fitted our site-specific models to the DMS data and sequence alignment, given the published tree for each dataset, and recorded the likelihoods (Supp. Table S1). We also identified the best-fitting site-homogeneous models given the alignment and tree topology using the Akaike Information Criterion and ProtTest software (Posada and Buckley 2004) and fitted these to the same alignment and tree. In all three cases, the site-specific models fit the data much better than the best site-homogeneous model (Table 1). This indicates that the site-specific models are better descriptors of the underlying substitution process. Extensive compositional heterogeneity is also apparent in the steady-state equilibrium frequencies of the fitted models, which differ dramatically among sites and when fit to different proteins in the same family (Fig. 2B, Supp. Figs. S4-S6).

**Table 1.** Fit of substitution models to sequence data.

| System/Model | Log-likelihood | AIC | ΔAIC | No. of parameters |
| --- | --- | --- | --- | --- |
| **SR** |  |  |  |  |
| SS-ER | -1689.7 | 3403.4 | 0 | 12 |
| SS-MR | -1768.2 | 3560.4 | 157.1 | 12 |
| JTT<sup>a</sup> | -1787.4 | 3614.8 | 211.4 | 20 |
| Poisson | -2042.0 | 4086.1 | 682.7 | 1 |
| **RBD** |  |  |  |  |
| SS-SARS | -1875.4 | 3774.9 | 0 | 12 |
| WAG<sup>a</sup> | -2090.4 | 4220.8 | 446.0 | 20 |
| Poisson | -2322.8 | 4647.6 | 872.7 | 1 |
| **HA** |  |  |  |  |
| SS-H1 | -21339.4 | 42702.8 | 0 | 12 |
| SS-H3 | -21516.0 | 43056.0 | 353.2 | 12 |
| FLU<sup>a</sup> | -21711.4 | 43462.8 | 760.0 | 20 |
| Poisson | -24825.8 | 49653.5 | 6950.7 | 1 |

All models include a $\Gamma$ 4 distribution. <sup>a</sup> Includes ML estimates of amino acid frequencies (+X)

### Reconstructed sequences are robust to unincorporated among-site heterogeneity

To characterize the robustness of ASR to incorporating or excluding compositional heterogeneity in the substitution model, we inferred ancestral sequences at every node in all three phylogenies using the site-specific model and the best-fitting site-homogeneous model for each dataset. For the SR and HA datasets, there are two DMS libraries in different backgrounds in each family and therefore two site-specific models: our initial assessment focuses on the model that fits the alignment best using AIC (Table 1). We also reconstructed ancestral sequences using a Poisson model, where all amino acid exchange rates are equal, as a reference case for a model that incorporates neither compositional heterogeneity nor any bias in the substitution process among amino acids. At each site in every ancestral node on the tree, we inferred the maximum a posteriori (MAP) state. To incorporate statistical uncertainty, we also inferred the set of plausible states, defined as the union of the set of MAP states at all sites and the set of alternative plausible states (all other states with posterior probability > 0.2) (Eick et al. 2017).

Despite the large differences in model fit, the inferred ancestral sequences are almost identical irrespective of the model used in all three datasets. Across all ancestors reconstructed in all three proteins, more than 98% of sites are reconstructed with the same MAP amino acid when the site-specific model, the site-homogeneous model, and even the Poisson model are used (Fig. 3A, lower diagonal). When statistical uncertainty is incorporated using the set of plausible states, >99% of sites have identical reconstructions across models (Fig. 3A, upper diagonal).

**Figure 3.**
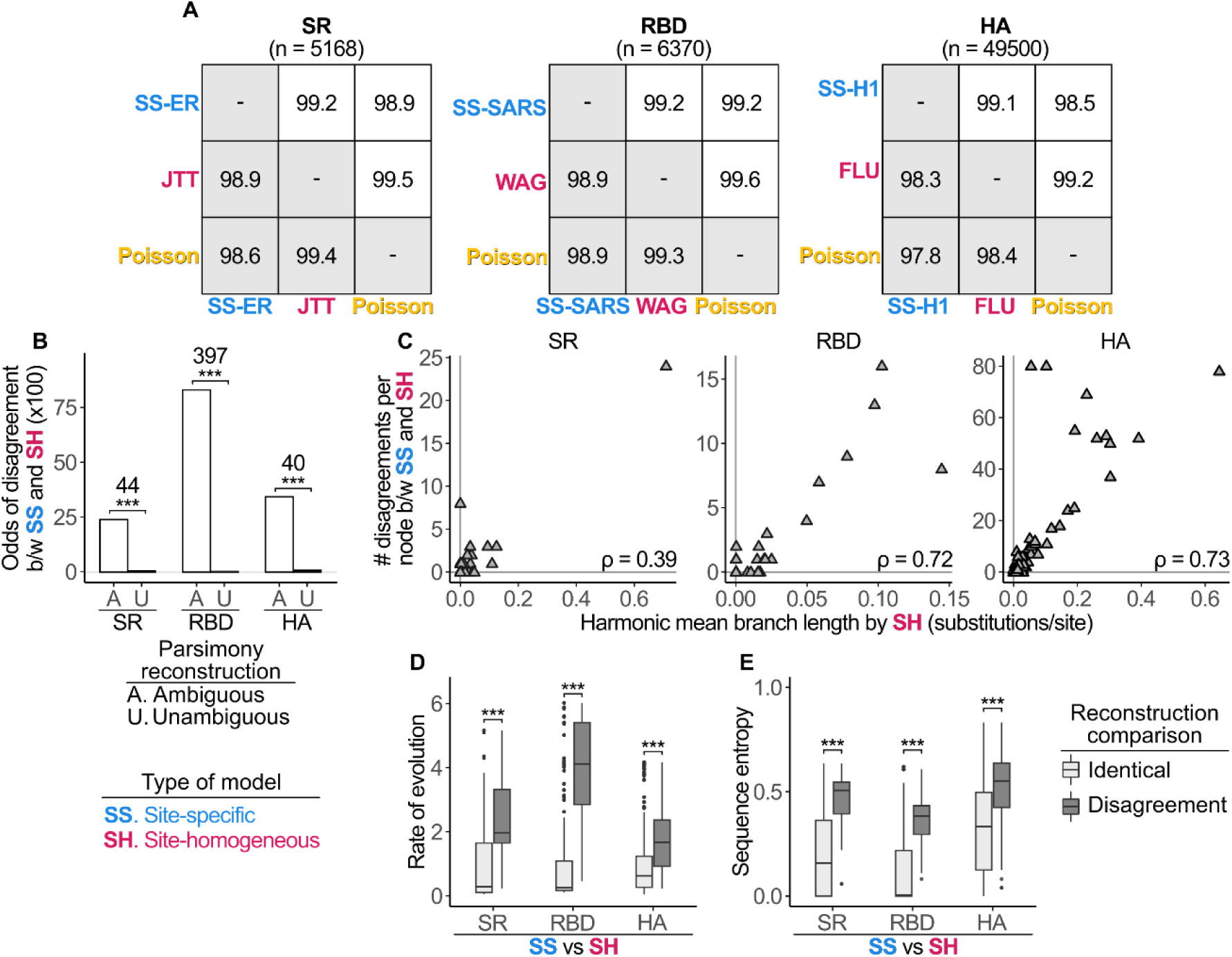
Robustness of ancestral sequences to unincorporated model heterogeneity due to strong phylogenetic signal. **(A)** Percentage of states across all nodes that are identically reconstructed states between models. *Lower diagonal*, identity between MAP states. *Upper diagonal*, identity when ambiguity is incorporated; the fraction of sites at which the MAP state using the model specified for that column is a member of the set of plausible states using the model for the row. Plausible states are defined as having PP > 0.2. *n*, total number of reconstructed sites in each protein system across all nodes. **(B)** Odds that the site-specific and site-homogeneous ancestral reconstructions disagree, for sites with maximum parsimony reconstructions that are is ambiguous (A) or unambiguous (U). Odds of disagreement in each category is defined as the number of reconstructed states that disagree between models divided by the number that agree. Odds ratios between categories are shown; ***, p < 0.001 by Fisher’s exact test for SR and RBD and chi-square test for HA (See Supp. Table S2). **(C)** Relationship between length of branches connected to a node and the number of disagreements between site-specific and site-homogeneous models at that node. Each point represents the reconstruction at one ancestral node. ρ, Spearman’s correlation coefficient. **(D-E)** The distribution of evolutionary rates **(D)** or sequence entropy **(E)** is shown for sites with identical reconstructions between site-specific and site-homogeneous models (light grey) and disagreeing reconstructions (dark grey). ***, p < 0.001 by a two-sample Kolmogorov-Smirnov test (See Supp. Table S4).

Incorporating compositional heterogeneity therefore has a minimal effect on ancestral reconstruction, indicating that ASR is largely robust to violation of the assumption of compositional homogeneity when site-homogeneous models are used. Of the few sites that differ among reconstructions, about half the differences are resolved by incorporating statistical uncertainty using the set of plausible states, indicating that, in most cases, the difference between the models at these sites is to prefer one plausible but uncertain state over the other.

### Phylogenetic signal is the primary determinant of ancestral sequence reconstruction

It seems surprising that the ancestral reconstructions are nearly identical across models, given that the site-specific model fits the data so much better, and compositional heterogeneity is rampant in the data (Fig. 2B). Why is ASR apparently robust to violation of the assumption of compositional heterogeneity? The Poisson model gives almost identical reconstructions, so ASR even appears to be robust to all forms of model heterogeneity (other than ASRV).

We reasoned that phylogenetic signal, not the substitution model used, is the primary determinant of ancestral sequence reconstruction. Phylogenetic signal is defined as the retention of the states found in an ancestral node in the nodes that are connected to it. Phylogenetic signal degrades as branches become long and sites become saturated with substitutions (Derrickson and Ricklefs 1988; Blomberg and Garland 2002). When phylogenetic signal is strong, there is a single most-parsimonious state, which minimizes the number of sequence changes on branches near the node being reconstructed; the only way in which different evolutionary models could yield different ancestral sequence reconstructions is if one of the models favors a state other than the parsimony state so strongly that multiple substitutions along the descendant branches become more probable than retaining the ancestral state. When phylogenetic signal is absent, however – when all neighboring nodes have different states – then several possible ancestral states imply the same number of substitutions on neighboring branches; biases in the model and the lengths of the neighboring branches will together determine which of these states is the maximum-parsiomony state. The model is therefore most likely to matter when phylogenetic signal is weak or absent.

If phylogenetic signal is the primary determinant of ancestral reconstruction, we expect that disagreements among models will happen predominantly at sequence sites and ancestral nodes where phylogenetic signal is absent. We evaluated this hypothesis using several measures of phylogenetic signal. First, we considered whether disagreements among models occur preferentially at sites where the maximum-parsimony reconstruction is ambiguous (Hillis and Huelsenbeck 1992). We performed parsimony-based ASR and classified every reconstructed site in each ancestor as unambiguous if and only if there is a single most-parsimonious ancestral state. We found that the disagreements between the site-specific and best site-homogeneous models are highly enriched at sites where the parsimony reconstruction is ambiguous compared to parsimony-unambiguous sites (Odds ratios between 40 and 400 for the three datasets, Fig. 3B, Supp. Table S2). Moreover, the sites that are ambiguously reconstructed using ML tend overwhelmingly to be associated with sites with ambiguous maximum-parsimony reconstructions (Odds ratios between 65 and 760, Supp. Table S3). These results corroborate the hypothesis that sites that retain phylogenetic signal will have consistently higher odds of being reconstructed identically irrespective of whether site-heterogeneity is included in the model.

A second indicator of phylogenetic signal is the amount of divergence between related sequences along a tree: Nodes connected by long branches will retain fewer ancestral states. For every node on each phylogeny, we calculated the harmonic mean distance of the three nodes it is connected to and then examined the association with the number of sites at that node where the site-specific and best site-homogeneous models disagree on the ancestral sequence state. For all three datasets, the number of disagreements was correlated with the harmonic mean distance (Fig. 3C). Distantly connected nodes are therefore more sensitive to incorporating compositional heterogeneity in the model.

The third indicator of the retention of phylogenetic signal is the evolutionary rate of a site, because fast-evolving sites will fail to retain ancestral states over shorter branches than slowly-evolving sites. We estimated the rate of evolution of each site as the posterior mean rate multiplier from the gamma distribution of ASRV. We found that the average rate of evolution is 3-16-fold higher at sites where the site-specific and site-homogeneous models disagree on the ancestral state than at the much larger number of sites where they agree (Fig. 3D, Supp. Table S4).

The final indicator of retention of phylogenetic signal is the conservation/variability of amino acids among extant sequences, measured as the Shannon entropy of the frequency distribution of amino acid states at each site in the alignment. An entropy of zero indicates that all extant sequences have the same state, the maximum possible retention of phylogenetic signal; sites with entropy of 1 have been completely randomized, with all 20 amino acids at equal frequency. As predicted, the entropy is much higher at sites where the site-specific and site-homogeneous models disagree than it is at the sites where they agree (Fig 3E, Supp. Table S4).

Taken together, these data indicate that phylogenetic signal, rather than the substitution model, is the primary determinant of inferred ancestral states; incorporating compositional heterogeneity into the model changes the inference only when phylogenetic signal is weak. These observations can also help explain why even when the site-specific and site-homogeneous reconstructions differ, the state preferred by one model is usually within the set of plausible states inferred by the other (Fig. 3A). Sites with reconstructions that are parsimony-ambiguous, due to a partial loss of phylogenetic signal, are typically ML-ambiguous, too (Supp. Table S3); in these cases, different models may prefer different states in the plausible set. Using different models – even incorporating compositional heterogeneity – rarely elevates a state that has very low likelihood under one model to having very high likelihood given another. When the site-specific and site-homogeneous reconstructions differ, the state preferred by one model is usually within the set of plausible states inferred by the other; partially retained phylogenetic signal results in a small set of plausible states, and the models choose differently among these options.

### ASR accuracy is unaffected by unincorporated among-site heterogeneity

Although ASR using different models yields identical reconstructions at most sites, it is possible that incorporating heterogeneity improves accuracy at the sites where they do disagree. Accuracy of ASR cannot be directly assessed using real proteins, because the true ancestral sequences are not known. We therefore examined how realistic forms of compositional heterogeneity affect the accuracy of ASR using phylogenetic simulations. For each of our three protein model systems, we simulated evolution of proteins of the same length as the real proteins, using the estimated parameters of the best-fitting site-specific model (Fig. 4A, Table 1, Supp. Table S1). To facilitate understanding how branch lengths – and the consequent loss of phylogenetic signal – affect ASR, we used a simple four-taxon tree, with equal branches of variable length from 0.05 to 1.6 substitutions per site, with 100 to 1000 replicates per branch length. We recorded the “true” ancestral sequences at both internal nodes of each replicate. We then inferred the ancestral sequences from the alignment of sequences at the tree’s tips using the true site-specific model or the best-fit site-homogeneous model, as well as the Poisson model; branch lengths and all free model parameters were optimized for each replicate alignment. Accuracy of ASR was measured as the percentage of sites at which the MAP reconstructed state matches the true ancestral state (Fig. 4B).

**Figure 4.**
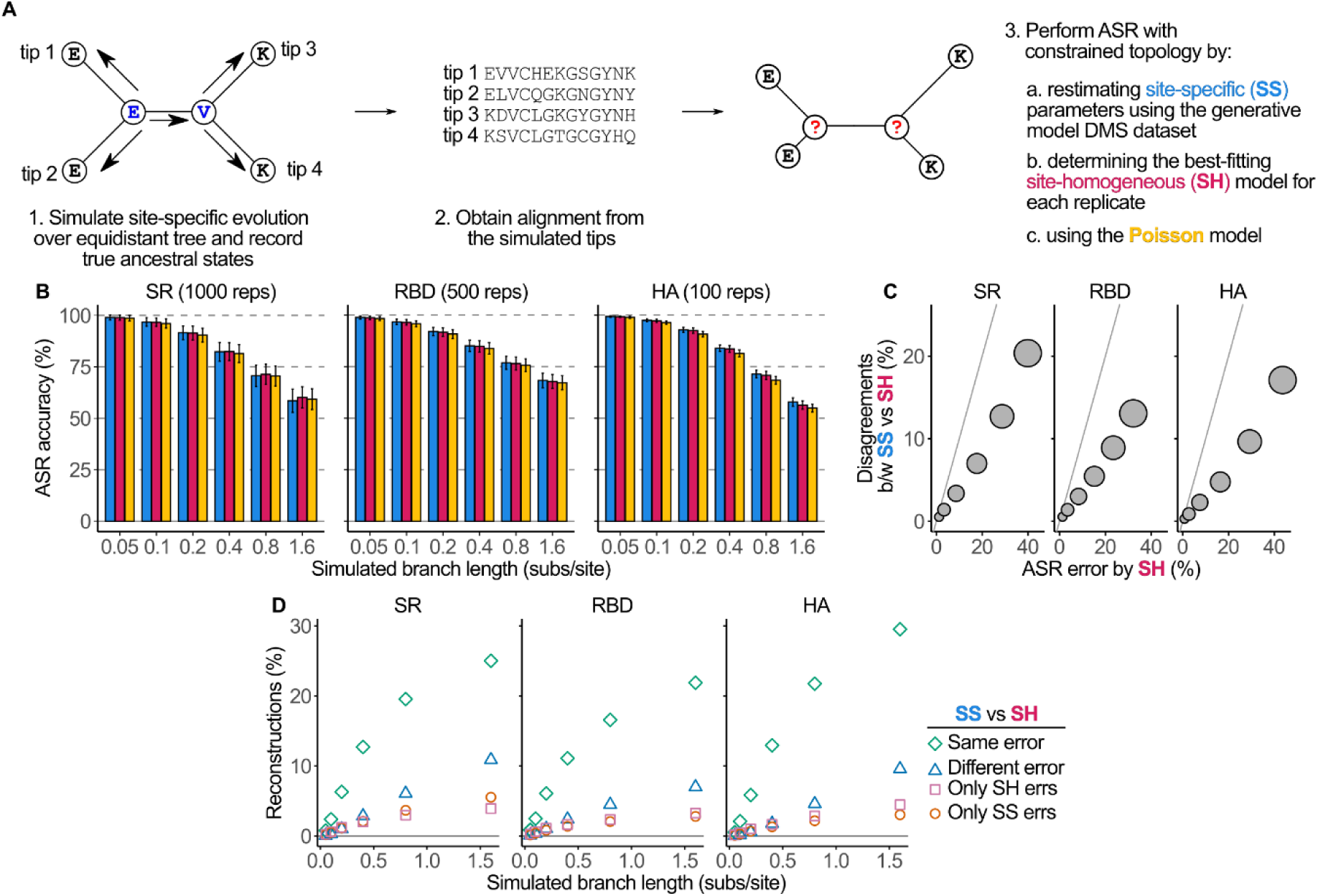
Reconstruction accuracy is mostly independent of model assumptions of site heterogeneity. **(A)** Workflow to simulate site-specific evolution and assess the accuracy of ancestral reconstruction. For each replicate, an ancestral sequence was seeded from the empirical site-specific model and allowed to evolve across the tree under that model. The trees have equal branch lengths, and simulations were performed across a range of lengths. The resulting alignment was then analyzed by maximum likelihood given the true topology and either the site-specific model, the best-fitting site-homogeneous model, or the Poisson model; branch lengths and free parameters were optimized by ML, ancestral sequences were inferred and compared to the true ancestral sequences. **(B)** Accuracy of ancestral reconstructions given the true site-specific model (blue), best site-homogeneous model (magenta), and Poisson (yellow) at all the simulated branch lengths. Column height shows the mean of replicates; error bars, standard deviation. The number of repetitions per system and per branch length is shown. **(C)** Reconstruction errors outnumber disagreements between models. Each circle represents one set of branch lengths and is plotted by the mean number of erroneous reconstructions by the site-homogeneous model and the mean percentage of disagreements between the site-specific and best site-homogeneous reconstructions across replicates. The size of each circle represents the branch length (from 0.05 to 1.6b substitutions per site). Gray line, y=x. **(D)** Percentage of reconstructions for which site-homogeneous and site-specific models assigned the same erroneous state (green), site-specific and site-homogeneous assigned different erroneous states (blue), and only site-specific (orange) or only site-homogeneous (pink) assigned an erroneous state.

The simulation results show that all three models have nearly identical accuracy (Fig. 4B, Supp. Table S5). At short branch lengths (≤0.2), all models yield highly accurate reconstructions and are nearly indistinguishable from each other. Accuracy declines as branch lengths increase irrespective of the model used, consistent with the finding that phylogenetic signal is the major determinant of inferred sequences. In two of the three proteins, the best site-specific model is slightly more accurate on average at longer branch lengths, but the difference is small. Even at the longest branch length – 1.6 substitutions per site on every branch – the difference in ASR accuracy between the site-specific model used to generate the data and the best-fitting site-homogeneous model is ∼1 percent point (Supp. Table S5). This corresponds to a total of 1 to 4 more errors per protein under the most challenging conditions for ASR. Even Poisson reconstructions are, at worst, less accurate than ancestral sequences inferred using the true model by ∼3 percent points.

The models could have similar accuracy because they infer the same sequences – and therefore make the same errors – or because they infer different sequences that contain a similar fraction of incorrect states. We compared the reconstructed sequences and found that the fraction of disagreements between site-specific and site-homogeneous models is relatively small, and there are more errors than disagreements (Fig. 4C), indicating that models often make the same error. To directly assess the relationship between errors and disagreements, we categorized reconstructions at each site and found that the majority of errors indeed involve both models choosing the same incorrect ancestral state at the same sites (Fig. 4D). The next most common outcome – accounting for no more than 10% of reconstructions, even under the conditions that produce the highest error rates – is for both models to err at a site but to choose different ancestral states. The least common outcome is for one model to get the reconstruction right and the other to get it wrong; the two models have similar error rates at these sites.

These results show that site-homogeneous reconstructions are virtually as accurate as site-specific reconstructions at a wide range of reconstruction conditions. Failure to incorporate realistic levels of heterogeneity into the substitution model therefore does not strongly affect the accuracy of the reconstructions, and it rarely affects the amino acid that is inferred.

### Phylogenetic signal determines the accuracy of ASR

Why do the models typically make the same error at the same sites? Because phylogenetic signal is the primary determinant of ancestral reconstruction, we hypothesized that the major source of error is misleading phylogenetic signal – convergence or reversal that makes the most parsimonious reconstruction incorrect. To test this hypothesis, we categorized sites by whether phylogenetic signal is honest (i.e., the maximum-parsimony state matches the true ancestral state), misleading, or ambiguous, and then examined the frequency of erroneous ML reconstructions in each category (Fig. 5A). At short branch lengths, virtually all parsimony reconstructions are true; as branch length increases, the proportion of sites with misleading or ambiguous signal increases, but even at the longest branch lengths, the majority of sites retain honest signal (Fig. 5B).

**Figure 5.**
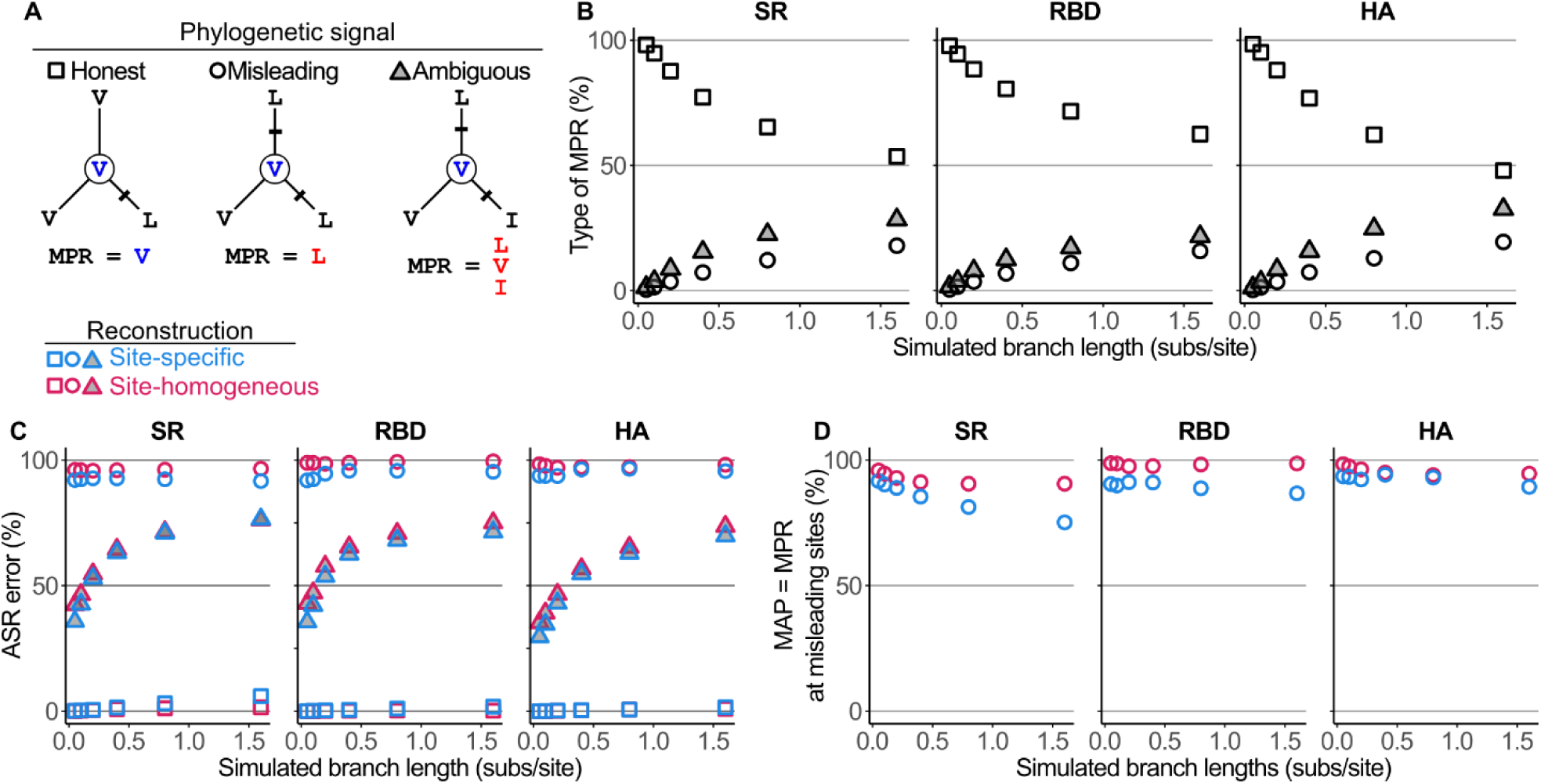
Misleading phylogenetic signal is the main cause of error in ASR. **(A)** Scheme for classifying phylogenetic signal by the correspondence of the maximum parsimony reconstruction (MPR) to the true ancestral state at a given site. Each subtree shows an example of an ancestral node with its true state (middle circle) and three connected nodes with their true states; the MPR given the states at the connected nodes is also shown. The phylogenetic signal of a site is *honest* if the MPR matches the true ancestral state because the state is retained in the majority of connected nodes (left); *misleading* if the MPR is different from the true ancestral state ancestor due to convergence/reversion (center); and ambiguous if there is no single MPR due to the lack of historical resolution (right). **(B)** Loss of phylogenetic signal and accumulation of misleading/ambiguous phylogenetic signal as a function of branch lengths. Each data point represents a set of replicate simulations, plotted by the branch length for the simulation and the percentage of MPRs that are honest (squares), misleading (circles), or ambiguous (triangles). **(C)** Percentage of errors in reconstructed ancestral states classified by their phylogenetic signal as described in panel **A.** Data from site-specific and site-homogeneous models are shown in blue and magenta, respectively. **(D)** Of reconstructions with misleading phylogenetic signal, the percentage at which the maximum a posteriori (MAP) state and the MPR are identical and erroneous is shown.

Our analysis corroborates the hypothesis that the primary source of error is misleading phylogenetic signal. When the signal is honest, the parsimony reconstruction is true, and the MAP state is almost always the same, irrespective of the branch lengths. When phylogenetic signal is misleading, the MAP state is almost always incorrect (Fig. 5C) and this occurs because the MAP state almost always matches the erroneous parsimony reconstruction (Fig. 5D). Errors also occur sometimes when phylogenetic signal is ambiguous, especially when branches are long, but to a lesser extent than when the signal is misleading (Fig. 5C).

The site-specific and -homogeneous models are similarly sensitive to the loss of honest phylogenetic signal. The site-specific model is very slightly less error-prone when phylogenetic signal is misleading but slightly less accurate when the signal is honest (Fig. 5C-D). Presumably, the reason why the site-specific models are more likely to choose a non-parsimonious state is that their biases for any particular amino acid states tend to be higher at multiple sites (Fig. 2B). Taken together, these data show that when branch lengths are short – making phylogenetic signal strong and honest – then ASR error is rare; when branch lengths are long, the most common cause of is convergence and reversal, not model misspecification. Neither the site-specific nor site-homogeneous models can overcome misleading phylogenetic signal.

### Model misspecification inflates confidence in ASR when phylogenetic signal is weak

When using ASR, we want to know both the best estimate of the ancestral state and also the degree of support for that site. The measure of confidence in reconstructed ancestral sequences is the posterior probability (PP), which expresses the probability that a state is correct, given the data, the model, and prior probabilities on amino acid frequencies and all other free parameters. Although the use of site-homogeneous models has a weak effect on the sequences inferred by ASR, it is possible that the use of an oversimplified model could inflate or deflate PPs, leading to over- or underestimated confidence in the MAP sequence compared to using a more realistic site-specific model.

To assess the effect of model violation on PPs, we binned the reconstructed states by their inferred PP using site-specific or site-homogeneous models and then computed the probability that the states in each bin are actually correct Fig. 6A). When branch lengths are short (0.05 to 0.2 substitutions/site), both site-specific and site-homogeneous models yield PPs that on average closely match the probability that the inferred state is correct in all three protein systems. As branch lengths become longer, however, site-homogeneous models yield PPs that overestimate the accuracy of MAP states; site-specific models yield PPs that are less inflated. Overestimation is most extreme at moderate levels of confidence: states with an inferred PP=1.0 are almost always correct, but when PP is between 0.50 and 0.95, the probability that the state is correct tends to be lower than the PP; this bias is much worse when site-homogeneous models are used than site-specific models (Fig. 6B, Supp. Fig. S7).

**Figure 6.**
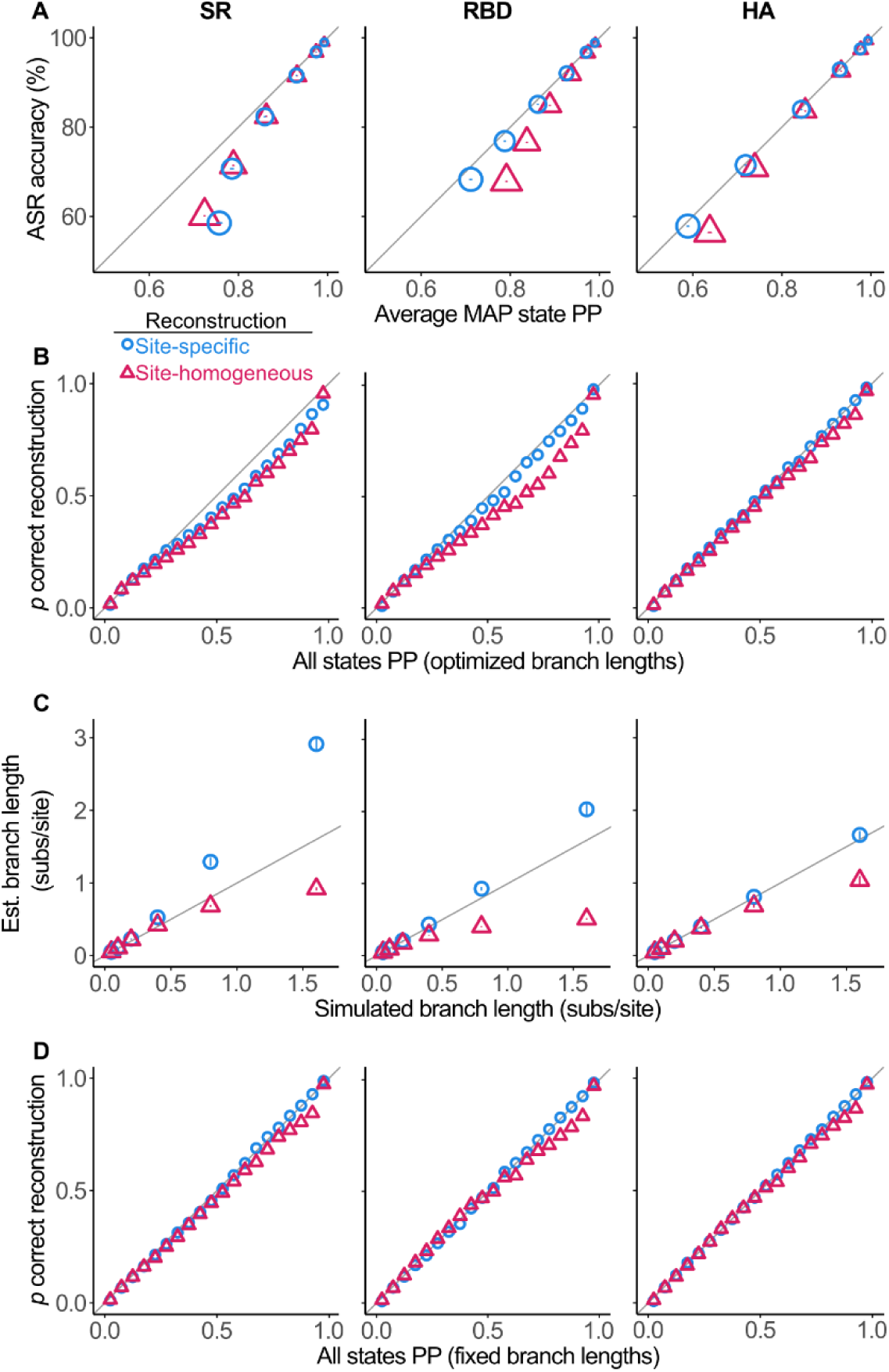
Posterior probabilities (PP) of ancestral states are inflated because of branch length estimation errors. Comparisons of mean ASR posterior probability versus the fraction of correct reconstructions from simulations given the true site-specific model (blue circles) or the best site-homogeneous model (magenta triangles) Gray line, y=x. **(A)** PPs are most inflated when branch lengths are long, irrespective of the model used. Each shape plots the mean PP of the MAP state and the fraction of correct reconstructions for replicate simulations under one set of branch length; shape size is proportional to the branch length (from 0.05 to 1.6 substitutions/site). Error bars, standard deviation of PP. **(B)** For reconstructions at branch length = 0.8, all possible ancestral states were binned by PP. Each shape plots the fraction of correct states among reconstructions in a PP bin. For other branch lengths, see Supp. Fig. S7. **(C)** Relationship of true branch lengths to the estimated branch length using each model; each shape plots the mean estimated branch length; error bars, standard deviation. **(D)** Same as (B) when branch lengths used for ASR are fixed to the true length.

Posterior probability values are conditioned, among other things, to the chosen substitution model and its parameters, as well as the tree and its branch lengths. The simulated data were generated using a site-specific model, so overestimation of confidence by site-homogeneous models could be caused directly by a mismatch between the model used for ASR and the true model; however, confidence was also overestimated when site-specific models were used for the SR and RBD simulations, even though these are the true generating models (Fig. 6A-B). We therefore hypothesized that inflated posterior probabilities were caused by differences between the inferred branch lengths and the true lengths. Consistent with this possibility, overconfidence is most severe when branch lengths are long, and it is under these conditions the branch length estimates are most inaccurate (Fig. 6C).

To directly test whether the inflation of PPs is caused indirectly by being conditioned on inaccurate estimates of branch lengths, we repeated ASR but this time fixing branch lengths to their true values used to generate the simulations. Under these conditions, overconfidence using site-homogeneous models is dramatically reduced; when site-specific models are used, the inferred PPs match the probability that an inference is true almost perfectly (Fig. 6D). Branch length misestimation is therefore the primary cause of overconfidence.

Taken together, these results demonstrate that PPs may overestimate the probability that an ancestral state is true when branch lengths are long, especially when site-homogeneous models, which tend to underestimate branch lengths, are used. This phenomenon is an indirect result of model violation: its primary cause lies not in its effect on the ASR calculation itself but in the estimation of branch lengths by oversimplified models, on which the ASR is conditioned.

### ASR is robust to unincorporated among-lineage compositional heterogeneity

We next investigated the effects of the second type of heterogeneity – changes in site-specific amino acid preferences among lineages caused by epistatic interactions with substitutions that occur as lineages diversify. To incorporate realistic levels of among-lineage heterogeneity, we compared robustness and accuracy when ASR uses different site-specific models parameterized by DMS experiments performed on distantly related homologous proteins within the same family, which have markedly different site-specific constraints (Fig. 7A).

**Figure 7.**
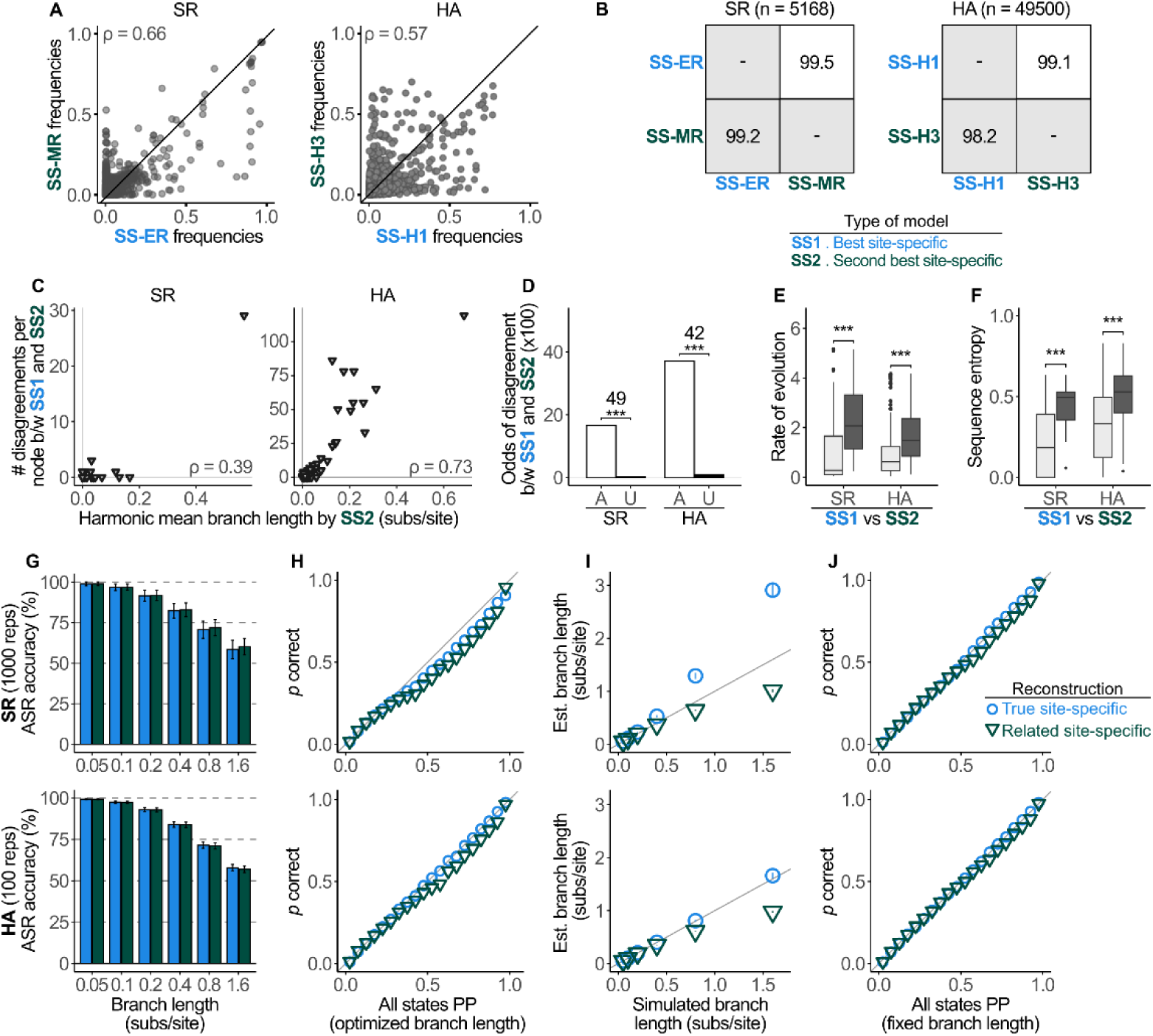
Among-lineage compositional heterogeneity does not affect ASR. **(A)** Comparison of the site-specific equilibrium frequencies between models fitted to experimental data from different proteins on the same phylogeny. Each point represents one amino acid state at one site in the protein, plotted by its equilibrium frequency for each of the two site-specific models. ρ, Spearman’s correlation coefficient. **(B)** Percentage of sites across all nodes with identically reconstructed states between models. *Lower diagonal*, identity between MAP states. *Upper diagonal*, identity when ambiguity is incorporated as in Fig 3A. **(C)** Relationship between length of branches connected to a node and the number of disagreements between site-specific models at that node. Each point represents the reconstruction at one ancestral node. ρ, Spearman’s correlation coefficient. **(D)** Odds that site-specific reconstructions disagree in sites where the parsimony reconstruction is ambiguous (A) or unambiguous (U). Odds ratios between categories are shown; ***, *p* < 0.001 by Fisher’s exact test for SR and chi-square test for HA (See Supp. Table S6). The distribution of evolutionary rates **(E)** or sequence entropy **(F)** is shown for sites with identical reconstructions between site-specific models (light grey) and disagreeing reconstructions (dark grey). ***, *p* < 0.001 by a two-sample Kolmogorov-Smirnov test (See Supp. Table S7). **(G)** Accuracy of ancestral reconstructions given the true site-specific model (blue) and the site-specific model parameterized using the closely related protein (green). Column height shows the mean of replicates; error bars, standard deviation. The number of repetitions per system and per branch length is shown. **(H)** For reconstructions at branch length = 0.8, all possible ancestral states were binned by PP. Each shape plots the fraction of correct states among reconstructions in a 5% bin. **(I)** Relationship of true branch lengths to the estimated branch length using each model; each shape plots the mean estimated branch length; error bars, standard deviation. **(J)** Same as (H) when branch lengths used for ASR are fixed to the true length.

In each family, the models for the homologous proteins differ substantially, indicated by the differences in their site-specific equilibrium frequencies (Fig. 7A, Supp. Figs. S4, S6). For SR, the steady-state equilibrium frequencies of amino acids at each site are moderately correlated between the ER and MR models (ρ = 0.66), with 71% of sites having the same most preferred state. For HA, H1 and H3 equilibrium frequencies show a slightly lower correlation (ρ = 0.57), and only 36% of sites with the same most preferred amino acid state.

Despite these differences, performing ASR with the models of homologous proteins yields almost identical results. When the empirical alignments are analyzed, the pairs of reconstructions are >98% identical in both SR and HA families; this identity increases to >99% when alternative plausible reconstructions are included (Fig. 7B). In both systems, the small number of disagreements again appears to be caused primarily by the loss of phylogenetic signal, with disagreements strongly enriched at sites and nodes with long branches, fast rates, high entropy, and parsimony-ambiguous reconstructions (Fig. 7C-F). These results suggest that realistic epistatic shifts in compositional heterogeneity have a very weak effect on ASR, changing the reconstruction only when phylogenetic information is lacking.

To understand how unincorporated among-lineage compositional heterogeneity affects the accuracy of ASR, we simulated sequence evolution as described in the previous sections using the site-specific model parameterized from the DMS experiment on one protein in the family, but now performed ASR using the site-specific model parameterized from the experiment on the homologous protein (Fig. 2B, Table 1). Accuracy using the pairs of models is indistinguishable, with no reduction in accuracy caused by using the model parameterized on a distantly related protein under any conditions examined (Fig. 7G).

Using the site-specific model from a distantly related homolog resulted in a weak overestimation of confidence, with PPs that exceed the probability that an inference is correct (Fig. 7H). This phenomenon again appears to be primarily attributable to underestimates of branch lengths, because fixing the branch lengths to their true values when performing ASR almost entirely eliminates the inflated confidence (Figs. 7I-J).

Taken together, these results indicate that model misspecification caused by among-lineage compositional heterogeneity has a minimal effect on the robustness or accuracy of ancestral sequence reconstructions. The functional effects of mutations at many sites does change as sequence substitutions accrue across the phylogeny, but these epistatic shifts have little effect on the inferred ancestral states. This insensitivity arises because phylogenetic signal, rather than differences in the models, is the primary determinant of the inference of ancestral sequence states.

## DISCUSSION

Our data show that ancestral sequences inferred by ASR are largely robust to realistic forms of unincorporated among-site and among-lineage compositional heterogeneity, provided that the alignments retain reasonable phylogenetic signal. When real sequence alignments from three protein families are analyzed with a site-specific model that incorporates compositional heterogeneity observed in functional experiments, the reconstructed sequences are nearly identical to those reconstructed using conventional site-homogeneous substitution models. When sequence evolution is simulated with experimentally derived forms of site-specific heterogeneity, using site-homogeneous models yields very similar reconstructions and nearly identical accuracy. The only exception is when branches are very long and phylogenetic signal is largely lost; in that case, accuracy remains virtually identical, but different models may make different errors. In addition, using site-specific models parameterized on distantly related protein family members also has a minimal effect on ASR sequences and accuracy; ASR is therefore also largely robust under realistic conditions to unincorporated among-lineage epistatic shifts in site-specific evolutionary constraints.

We do not claim that the site-specific models we derived from experimental data represent the “true” evolutionary model. Rather, our purpose was to incorporate into ASR a reasonable approximation of among-site heterogeneity in functional constraints and then compare the results to ASR using models that exclude this compositional heterogeneity. The DMS experiments on which our models are based measured the effects of mutations on the major biochemical or cellular functions of the proteins. Although the exact quantitative mapping of these functions to fitness is unknown, the logistic relationship in our model roughly approximates the effect of purifying selection against mutations at each site that impair a protein’s function. This information is entirely lacking from site-homogeneous models, which assume that sites in the protein’s core, on its surface, in its active site, and on unstructured loops are all subject to the same constraints. We found that incorporating empirically-derived heterogeneity using this kind of site-specific model dramatically improves the statistical fit to sequence alignments in all three protein families that we studied, but it barely changes ancestral sequence reconstructions or their accuracy. Given these findings, it seems unlikely that, if we precisely knew the among-site differences in selective constraints that pertained during history, incorporating them would strongly affect ASR or its accuracy.

Our findings are consistent with prior work in this area. A recent study found that sequences inferred by ASR and their accuracy are very similar irrespective of the particular site-homogeneous model used (less than 1% percentage point difference even when sequences differ at 50% of sites) (Del Amparo and Arenas 2022). Another study developed a biophysical model to account for site-specific effects on protein stability (SSS); when sequences were simulated under the SSS model and ASR was then performed, accuracy was nearly identical irrespective of whether the model used for ASR was the true SSS generating model or a variety of site-homogeneous models (Arenas et al. 2017).

We found that phylogenetic signal, rather than the substitution model, is the primary determinant of ancestral sequence reconstruction. When phylogenetic signal is present, the most parsimonious reconstruction is almost always assigned as the maximum a posteriori (MAP) state, irrespective of the model used. Different models yield different reconstruction only when phylogenetic signal is weak – when branches are very long and/or rates at a site are very high. These observations can be easily understood by considering how phylogenetic signal affects the probability of ancestral state reconstructions. Consider a node with strong phylogenetic signal, where all three neighboring nodes of a focal ancestral node share the ancestral state. The likelihood of the maximum-parsimony state at the focal node is the product of the likelihoods that the state will not change along each connecting branch. With an unbiased model, the likelihood of no-change on a branch will always be greater than that of a change to a particular different state, so the product of the no-change likelihoods will be greater than the product of the likelihoods of three convergent changes -- much greater if the branches are of short or moderate length. The only way for the MAP state to differ from the maximum-parsimony state is if the model is so biased that it overcomes this difference. By contrast, when phylogenetic signal is absent – that is, all neighboring nodes have different states – then any reconstruction requires at least two changes on the descendant branches, and which one of these has the highest posterior probability depends entirely on branch lengths and biases in the model. For different models to yield different reconstructions in this latter case, they must be biased towards different states. The model therefore affects the reconstruction mostly when there is no phylogenetic signal; when signal is present, different models yield different reconstructions only when branch lengths are long and the models are biased in favor of different states.

Our analyses provide insight into the causes of error in ASR. Using site-specific vs. site-homogeneous models has a negligible effect on accuracy; even the completely uninformative Poisson model performs almost as well. The loss of honest phylogenetic signal, not model misspecification, is the primary cause of erroneous reconstructions. When phylogenetic signal is strong, reconstructions are accurate regardless of model choice. When phylogenetic signal is misleading because of convergence or reversal, reconstructions are inaccurate regardless of model choice. When phylogenetic signal is ambiguous, some reconstructions err stochastically; the state inferred can be sensitive to model choice but no model is systematically more accurate than others.

Our findings have implications for the practice of ASR. First, the fact that site-specific models weakly affect ASR and its accuracy means that little will be gained by developing and using such models for the purpose of ASR. There is no need to perform laborious DMS experiments to parameterize empirical site-specific models. Even fitting heterogeneous models like the CAT model to sequence alignments, which is computationally costly, is unlikely to be beneficial for ASR per se; however, such site-specific models may have benefits for inferring topologies and estimating branch lengths (Lartillot and Philippe 2004; Si Quang et al. 2008; Schrempf et al. 2020; Szánthó et al. 2023). Second, our finding that epistatic shifts in compositional heterogeneity across a tree have a negligible effect on ASR and its accuracy means that there is no apparent need for ASR to incorporate these interactions into complex models, despite the fact that epistatic interactions are widespread in the genetic architecture of proteins (Breen et al. 2012; Starr and Thornton 2016), and epistatic shifts in evolutionary rates can result in incorrect inference of tree topologies (Kolaczkowski and Thornton 2004; Philippe et al. 2005; Spencer et al. 2005).

Instead of using complicated models, the most effective way to improve the accuracy of ASR in practice is to maximize phylogenetic signal by densely sampling sequences around the nodes of interest. Our data show that this strategy will not only reduce the number of model disagreements in ASR and also increase reconstruction accuracy; previous work establishes that this is also an effective strategy for improving phylogenetic inference (Hillis 1996; Hillis 1998; Zwickl and Hillis 2002). Our data also show that shortening long branches also improves the correspondence between the calculated posterior probability of an ancestral state and the probability that the state is true. There may be cases in which this strategy cannot be implemented: in some protein families, there may be no extant sequences that can break up long branches near nodes of interest. If phylogenetic signal is irreparably weak, ASR should be approached with caution or not performed at all.

Although the misspecification caused by using site-homogeneous models has little overall effect on ASR and its accuracy, we found that the model used does changes the reconstruction at a small fraction of sites. Our data show that most of this model-related ambiguity in ancestral states can be incorporated by using the AltAll strategy for addressing statistical uncertainty in ASR (Eick et al. 2017). That is, most ambiguity caused by model misspecification overlaps with stochastic ambiguity; this is expected, because the model matters most when phylogenetic signal is weak -- the same condition under which statistical uncertainty is greatest (Eick et al. 2017). We therefore recommend that the AltAll strategy be used to incorporate both forms of uncertainty. This strategy, however, identifies ambiguously reconstructed sites based on their posterior probabilities, which can be overestimated when oversimplified models are used and branches are long, as our data and a recent study show (Sennett and Theobald 2023). Special caution should therefore be when reconstructing nodes with weak signal, because some states that should be considered ambiguous could be excluded.

Although reconstructed sequences are largely identical among models, model violation does change the inferred sequence state at a small fraction of sites. The goal of ASR is typically not to infer the exact ancestral sequence but rather to assess the protein’s biochemical and functional properties. In this study, we did not experimentally characterize the sequences reconstructed using site-specific models, so we cannot rule out the possibility that the small number of differences among reconstructed sequences could affect these properties. We predict, however, that disagreements between models are unlikely to strongly affect function. Several lines of evidence support this view. First, disagreements are very strongly enriched at fast-evolving, high-entropy sites with weak phylogenetic signal; these patterns occur primarily at sites that are subject to weak constraints, with states that are repeatedly sampled during evolution because they have indistinguishable effects on function. Second, experiments show that AltAll reconstructions, which incorporate most of the differences in ASR caused by using different models, generally have the same functional properties when assessed experimentally as the MAP reconstructions (Eick et al. 2017). Third, ancestral coral fluorescent proteins reconstructed using different models have almost identical fluorescence spectra when assessed experimentally (Ugalde et al. 2004). Finally, a recent study found that the predicted stability of proteins reconstructed using site-specific models differed from those reconstructed using site-heterogeneous models by only a few thousands to a few hundredths of a kcal/mol (Arenas et al. 2017). ASR errors caused by model violation can change the biochemical properties of the inferred residue, however, so it is possible that these errors may sometimes affect a protein’s functions (Sennett and Theobald 2023).

Although our findings are reassuring, we emphasize that ASR is by no means foolproof. Here we addressed two major forms of complexity that are rarely incorporated into the models used for ASR – among-site and among-lineage compositional heterogeneity – and found that this form of model violation does not reduce the accuracy of reconstructed sequences. But some errors in ASR are inevitable, especially when phylogenetic signal is weak. ASR should therefore be used on protein families in which the descendants retain phylogenetic signal of the sequence states that existed in their ancestors. The best way to improve accuracy is to curate alignments in which that signal is as rich as possible.

## METHODS

### Data and code availability

Data and code for analysis of the empirical datasets as well as summarized tables of the simulated data are available in github.com/JoeThorntonLab/site-specific-asr. Examples of ten replicates of simulated RBD evolution at 0.8 substitutions/site were included along with the scripts used to summarize data. ChatGPT-4o was used to optimize the performance and memory usage of all data analysis scripts used in this manuscript.

### Experimentally informed site-specific model fitting

The site-specific probabilistic model incorporating information from DMS functional experiments is based on the approach developed by Bloom (Bloom 2014b; Bloom 2014a). The key differences are 1) Bloom’s model used experimental measurements of fitness effects of mutations to estimate the probability that a mutation will be fixed, whereas our model uses measured effects on function and a simple function-to-fitness transformation; and 2) Bloom’s model estimated nucleotide mutation rates from nucleotide-based alignment data, whereas our model uses a general time-reversible model of nucleotide substitution and the genetic code to estimate the rate of producing amino acid changes by mutation from amino acid-based alignment data.

The model consists of a matrix, *q*, at each site *a*, which specifies for that site the instantaneous rates at which each of the twenty amino acids *x* is substituted to the 19 other amino acids *y.* Each rate *q_a,xy_* is the product of the rate at which an amino acid replacement is produced by mutation (*μ_xy_*) times the probability that the mutation will be fixed (*f_a,xy_*) (Halpern and Bruno 1998):

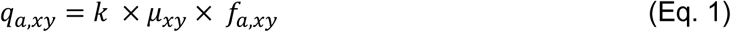

*k* is a normalization factor which scales the total rate of substitution so that the length of any branch equals the expected number of substitutions per site on that branch. The mutation rates do not vary among protein sites, whereas the fixation probabilities do, thus incorporating site-specific differences in the functional/fitness effects of each amino acid mutation.

#### Mutation rates

Each mutation rate *μ_xy_* from amino acid *x* to amino acid *y* can be decomposed into the equilibrium frequency of amino acid *y* (*π_xy_*) and the exchangeability from *x* to *y* (*ρ_xy_*):

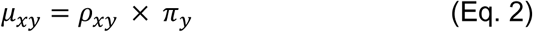

With 20 amino acids, a time-reversible version of this model would require 209 free parameters – 190 for the exchangeabilities and 19 for the equilibrium frequencies. The problem can be simplified by specifying the amino acid model in terms of the four possible nucleotides and the genetic code that maps DNA states to amino acids. We adopt this approach and use the general time-reversible model of nucleotide mutation and the standard genetic code. The equilibrium frequency *π_y_* of any amino acid *y* is the sum of the frequencies of all the codons that code for *y,* denoted as *codon(y):*

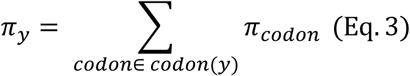

The frequency of a codon, denoted *αβγ* to specify the nucleotide at each of the codon’s three positions, is the product of the frequencies of the three nucleotides in the general time-reversible model:

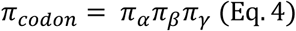

The exchangeability *π_xy_* between any two amino acids *x* and *y* is calculated from the exchangeabilities for single-nucleotide changes that can cause a codon for *x* to become a codon for *y.* Consider a change from a particular codon for *x* that contains nucleotide a (*codon x*_a_) to a codon for *y* that contains nucleotide b (*codon y*_b_). The exchangeability for that codon change is the exchangeability of the nucleotide mutation:

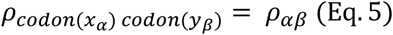

The total exchangeability from amino acid *x* to amino acid *y* is the sum of the exchageabilities for all single-nucleotide changes from codons for *x* to codons for *y,* each weighted by the relative frequency of codon *x*_a_ among all codons for *x:*

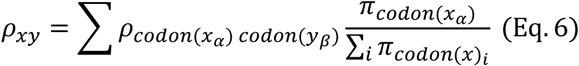

The rate of mutation between codons that require more than 1 nucleotide substitution is fixed at zero.

#### Site-specific probability of fixation

As in Bloom’s model, the probability that a mutation will be fixed is calculated from its selection coefficient, using the Kimura equation for a diploid population of size *N:*

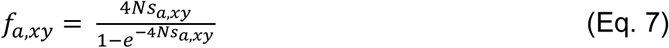

where s_a,xy_ represents the selection coefficient for the replacement of *x* by *y* (Kimura 1962; McCandlish and Stoltzfus 2014) (*eq. 5*). *N* determines the extent to which substitutions implied on the phylogeny are attributed to fitness differences or drift.

In Bloom’s, model, the fitness effect of each amino acid replacement at each site was measured directly in a DMS experiment, and selection coefficients could be directly calculated from them. In our datasets, protein function was measured, so we inferred fitness effects and selection coefficients from functional measurements using a simple function-to-fitness transformation. We use a simple sigmoid representation of purifying selection, which imposes an upper bound fitness (above which increases in function cause no further improvement in fitness) and a lower bound (below which reductions in function cause no further fitness decrement). Specifically, the growth rate *r_a,x_* of a genotype carrying amino acid state *x* at site *a* is related to the experimentally measured functional value *F_a,x_* for that genotype using the sigmoid relationship:

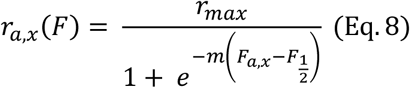

where *r_max_* is the upper-bound growth rate, *F_1/2_* is the midpoint (the function associated with half-maximal growth), and *m* determines the slope of transition from lower to upper bound. These free parameters apply to all states and all sites.

These inferred growth rates allow selection coefficients to be calculated for any substitution from amino acid *x* to *y* at site *a*:

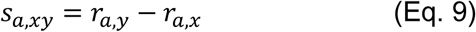

Altogether, the substitution model contains the 8 free parameters of the general time-reversible nucleotide mutation model, the slope and midpoint of the function-to-fitness transformation, and the selection tuning parameter *Nr_max_*11. (When equations 6 and 7 are substituted into equation 5, the product Ns in eq. 5 is formulated in terms of N r_max_, so this product can be treated as a single parameter.) These free parameters, along with the branch lengths, are estimated from the data by maximum likelihood as part of the phylogenetic optimization process; they take on the values that maximize the probability of observing all the sequence data, given the observed functional effects of mutations, the genetic code, and the tree topology.

#### Site-specific equilibrium frequencies

For the site-specific models, the steady-state equilibrium frequencies of each amino acid at every site (Φ*_a,y_*) serve as the priors for empirical Bayesian ancestral reconstruction (Yang et al. 1995). They can be derived from the model as follows:

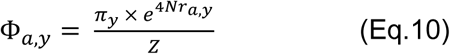

Where Z is a normalization factor that ensures that the frequencies of all 20 states at a site sum to one (∑*_y_* Φ*_a,y_* = 1). Logo plots of site-specific equilibrium frequencies were generated using *ggseqlogo* (Wagih 2017).

We also included among-site rate heterogeneity using a four-category discretized gamma distribution (Yang 1996), which requires estimation of a single additional free parameter (the shape parameter of the gamma distribution).

This model was implemented in Python as *DMSPhyloAA* and is publicly available at github.com/JoeThorntonLab/site-specific-asr. *DMSPhyloAA* can also perform ML optimization and ancestral reconstruction using a provided site-homogeneous model.

### Empirical datasets, model selection, ancestral sequence reconstruction and analysis

Alignments, phylogenetic trees, and DMS datasets were all obtained from previous publications. Sites with >50% gaps were removed from the alignments and their corresponding positions in the DMS datasets were eliminated. Missing data in the DMS datasets were considered to have no functional effect relative to the WT state at the corresponding site. The tree and alignment obtained from Park et al. (2022) were reduced to include sequences only from the steroid receptor family. The HA nucleotide alignment obtained from Hilton & Bloom (2018) was translated into amino acid sequences while maintaining the position of alignment gaps.

For each dataset, the best-fitting site-homogeneous model given the published topology was identified using *ProtTest* 3.4.2 and the corrected Akaike Information Criterion; all available models (JTT, LG, DCMut, MtREV, MtMam, MtArt, Dayhoff, WAG, RtREV, CpREV, Blosum62, VT, HIVb, HIVw, FLU) were compared (Guindon and Gascuel 2003; Darriba et al. 2011). Equilibrium frequencies for the site-homogeneous models were estimated from the data by maximum likelihood (+X) with *RAxML* 8.2 (Stamatakis 2014). The four-category discretized gamma distribution was also incorporated with site-homogeneous models (Yang 1996).

All ancestral sequence reconstructions were performed using *DMSPhyloAA,* given the published topology, while optimizing branch lengths and free model parameters. For alignment sites containing gaps in some sequences, maximum parsimony was used to determine whether a gap or sequence state should be present or absent. If a state is inferred to be present, the particular state was reconstructed by ML; if a state is inferred to be a gap, it is reconstructed as a gap irrespective of the model used.

For the analysis of phylogenetic signal, parsimony-based ASR was performed using *Mesquite* 3.70 (Maddison and Maddison 2021). The reconstruction of a site at a node was classified as parsimony-ambiguous if there are multiple equally parsimnonious states, and unambiguous if there is a single MP state.

The average distance of node to its immediate neighbors was calculated as the harmonic mean of the branch lengths coming off it. The harmonic mean reduces the effect of outliers, thus providing a conservative measurement of distance.

The rate of evolution per site under each site-homogeneous model was estimated using *PhyML* 3.0 (Guindon and Gascuel 2003; Guindon et al. 2010). The alignment Shannon entropy was calculated using the amino acid frequency per site using a logarithm base 21 (20 amino acid states + gaps).

### Site-specific sequence evolution simulation, ancestral sequence reconstruction, and analyses

To simulate sequences under site-specific models, we implemented a script, *simulate_alignment*, which uses the *simSeq* function of the R package *phangorn* 2.11.1 (Schliep 2011) Using the fitted site-specific model parameters for each dataset, we simulated sequence evolution on four-taxa trees with equally sized branch lengths. (For datasets with DMS experiments in multiple backgrounds, we used the model parameterized by the DMS experiment that yields the highest likelihood across the entire alignment/phylogeny.) The number of replicate simulations per site-specific model was varied because *DMSPhyloAA* processing time depends strongly on sequence length.

Ancestral sequence reconstruction was then performed given each simulated alignment, using the true generating tree topology. Site-specific reconstructions were obtained with *DMSPhyloAA.* For site-homogeneous reconstructions, the best-fitting site-homogeneous model for each alignment was identified using *ProtTest* 3.4.2 , empirical frequencies were estimated using *RAxML* 8.2 (Stamatakis 2014), and *PAML* 4.9j (Yang 1997; Yang 2007) was used to estimate branch lengths and reconstruct ancestral sequences. Parsimony-based ancestral reconstructions were obtained using the *ancestral.pars* function of *phangorn* 2.11.1 (Schliep 2011).

Posterior probability accuracy was evaluated by binning every possible reconstructed state at each node by their PP in 5% increments and calculating the fraction of true ancestral states in each bin. Bins with <200 elements were discarded. The effect of branch length optimization on posterior probability estimation was assessed by fixing branch lengths to their true values during ASR on an independent set of simulations made at 0.8 substitutions/site.

## ACKNOWLEDGEMENTS

We thank members of the Thornton Lab for comments and advice throughout the project. We thank Brian P.H. Metzger for contributions to conceiving the project and the DMS-site-specific model. This work was completed in part with resources provided by the University of Chicago’s Research Computing Center. This work was supported by National Institutes of Health grants R35-GM145336, R01-GM131128, R01-GM121931 (J.W.T.), and a Samsung Graduate Fellowship (YP).

