## Supplementary materials for "Robustness of ancestral sequence reconstruction to among-site evolutionary heterogeneity and epistasis"

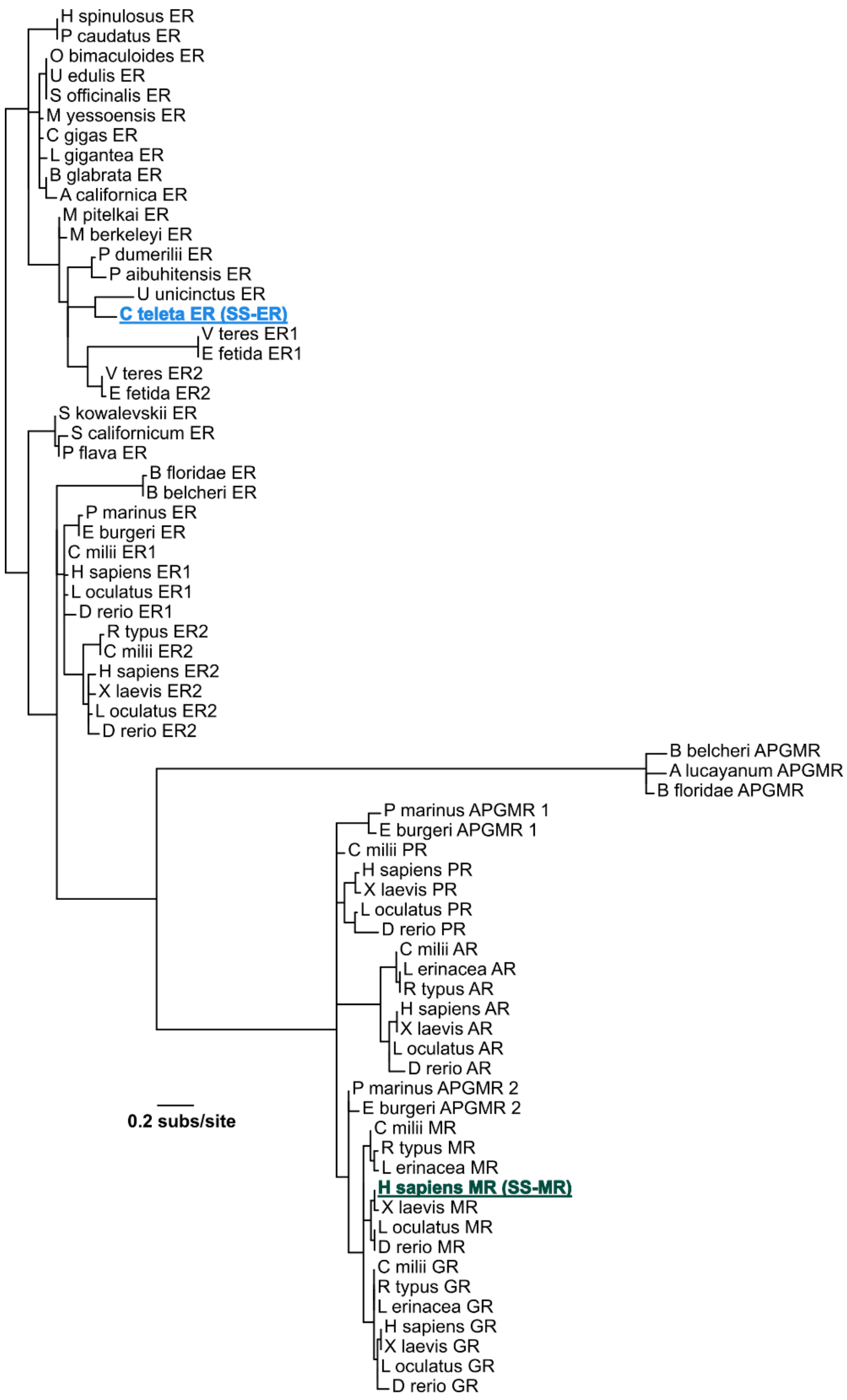


**Supplementary Figure S1. Maximum likelihood (ML) phylogeny of the DNA-binding domain of the steroid hormone receptor (SR).** Topology used for the analysis of SR data obtained from Park et al. (2022). Branch lengths were optimized using the model JTT+Γ4+X. The site-specific (SS) substitution models SS-ER and SS-MR were informed from the deep mutational scanning (DMS) data obtained from the background of *Capitella teleta* estrogen receptor (blue) and *Homo sapiens* mineralocorticoid receptor (green) DNA-binding domains, respectively.

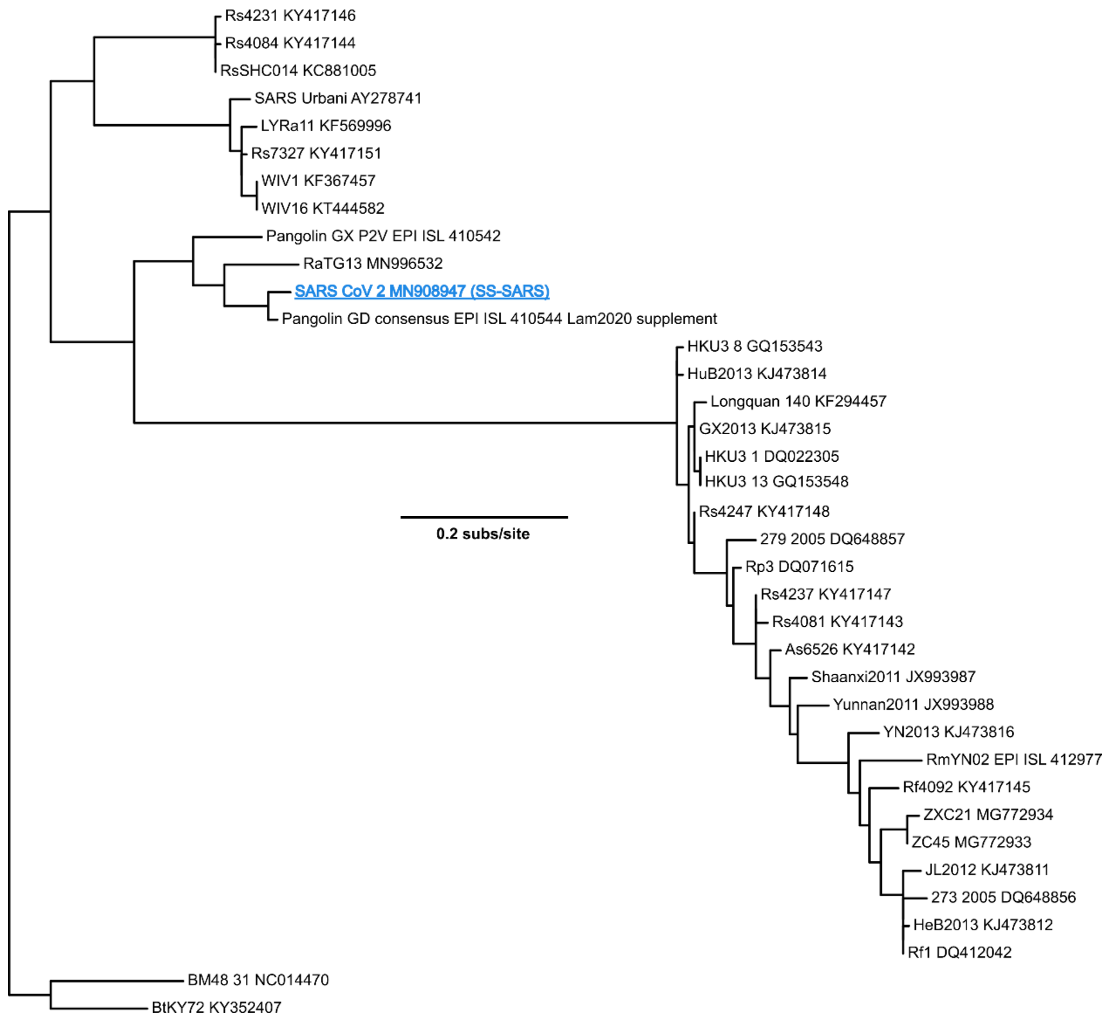


**Supplementary Figure S2. ML phylogeny of the spike glycoprotein receptor-binding domain (RBD).** Topology used for the analysis of RBD data obtained from Starr et al. (2020). Branch lengths were optimized using WAG+Γ4+X. The site-specific substitution model, SS-SARS, was informed by the DMS data on the background of SARS-CoV-2 RBD (blue).

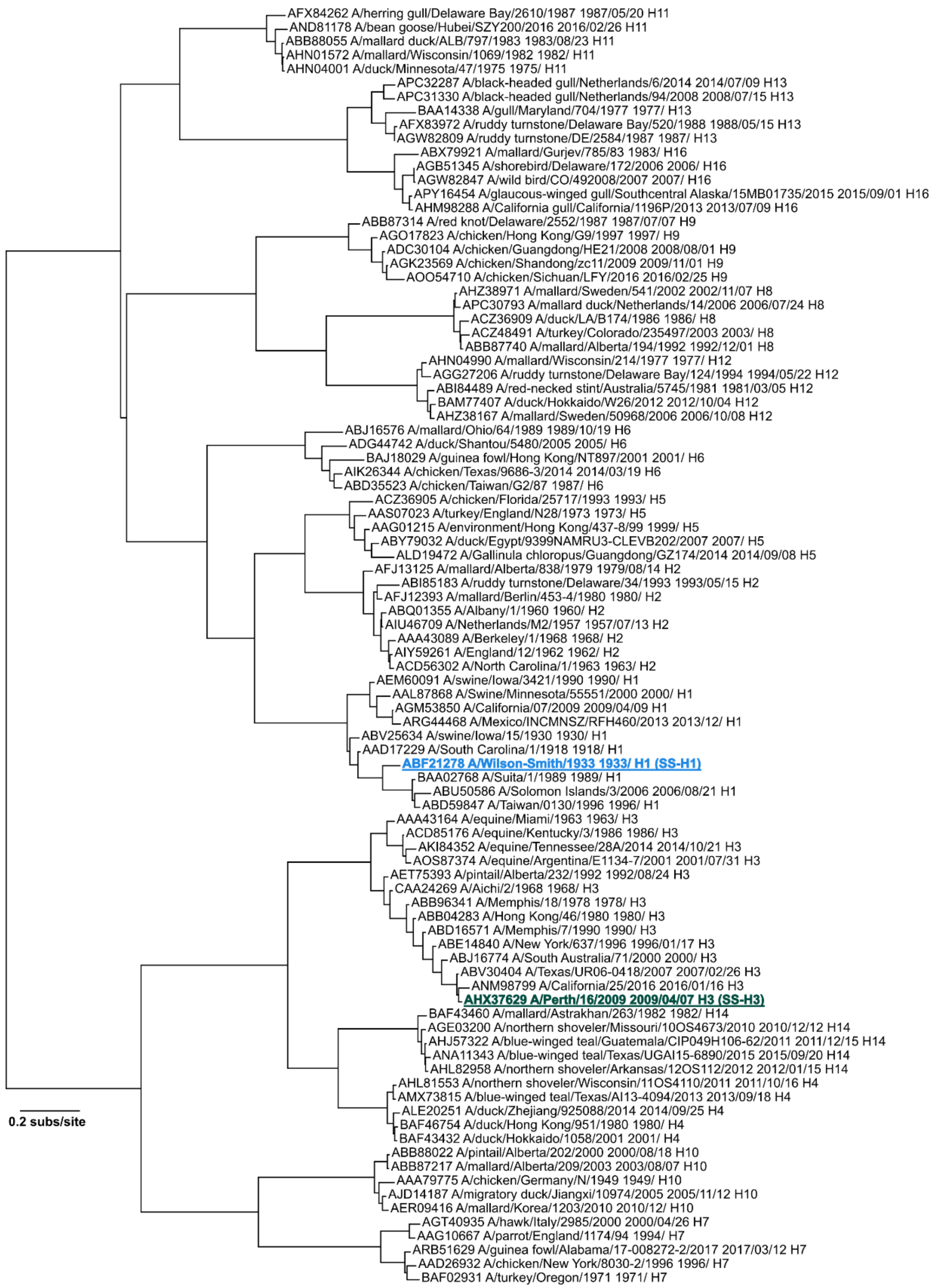


**Supplementary Figure S3. ML phylogeny of influenza A virus hemagglutinin (HA) subtypes.** Topology used for the analysis of HA data obtained from Hilton and Bloom (2018). Branch lengths were optimized using FLU+Γ4+X. The site-specific models SS-H1 and SS-H3 models were informed by the DMS data on the background of the HA subtypes H1 (blue) and H3 (green), respectively.

**Supplementary Table S1. Maximum likelihood estimates of parameters of site-specific models.**

| | SR | | RBD | HA | |
|---|---|---|---|---|---|
| Parameters | SS-ER[c] | SS-MR | SS-SARS[c] | SS-H1[c] | SS-H3 |
| Exchangeabilities[a] | | | | | |
| GA | 1.560 | 1.672 | 1.339 | 2.187 | 2.149 |
| GC | 0.602 | 0.608 | 0.716 | 0.618 | 0.672 |
| TA | 1.004 | 0.828 | 0.883 | 0.521 | 0.395 |
| TC | 1.730 | 1.652 | 0.750 | 1.476 | 1.282 |
| TG | 0.533 | 0.487 | 0.291 | 0.660 | 0.510 |
| Equilibrium frequencies[b] | | | | | |
| C | 0.174 | 0.167 | 0.204 | 0.181 | 0.166 |
| G | 0.266 | 0.265 | 0.198 | 0.216 | 0.223 |
| T | 0.249 | 0.246 | 0.297 | 0.214 | 0.239 |
| Logistic curve | | | | | |
| Max value ($Nr_{max}$) | 5.449 | 4.110 | 1.882 | 2.853 | 2.071 |
| Steepness (m) | 2.310 | 2.850 | 87.997 | 0.963 | 1.754 |
| Midpoint ($F_{½}$) | -1.218 | -1.127 | $-3.508 \times 10^{-2}$ | -1.592 | -0.961 |
| Gamma distribution | | | | | |
| Alpha | 1.215 | 0.833 | 0.567 | 1.325 | 1.177 |

Maximum likelihood estimates of the experimentally informed site-specific models per DMS dataset

[a] Unscaled values. The exchangeability CA is set to 1 in all cases.

[b] The frequency of A is calculated as 1-(C+G+T)

[c] Inferred parameters used to generate the site-specific simulations for each system

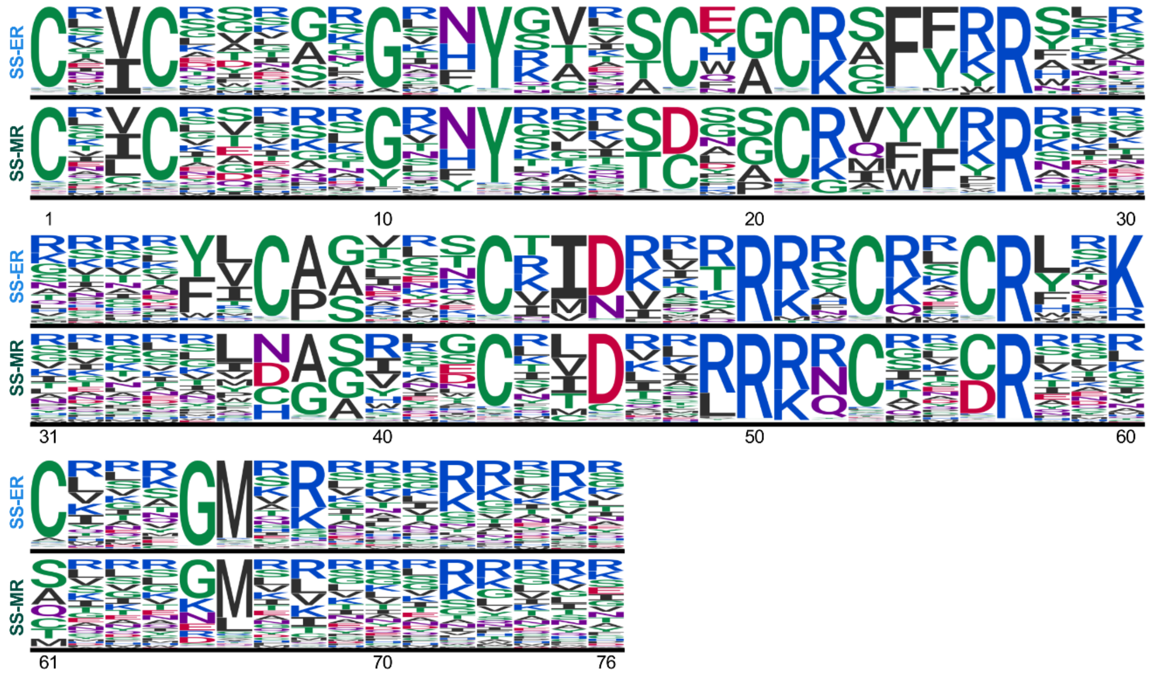


**Supplementary Figure S4. Compositional heterogeneity of the site-specific models of the steroid receptor.** Site-by-site comparison of the site-specific equilibrium frequencies estimated by the SS-ER (top) and SS-MR (bottom) models. At each site, states are organized from most to least frequent from top to bottom. Logo plots were generated using the R package *ggseqlogo* (Wagih 2017).

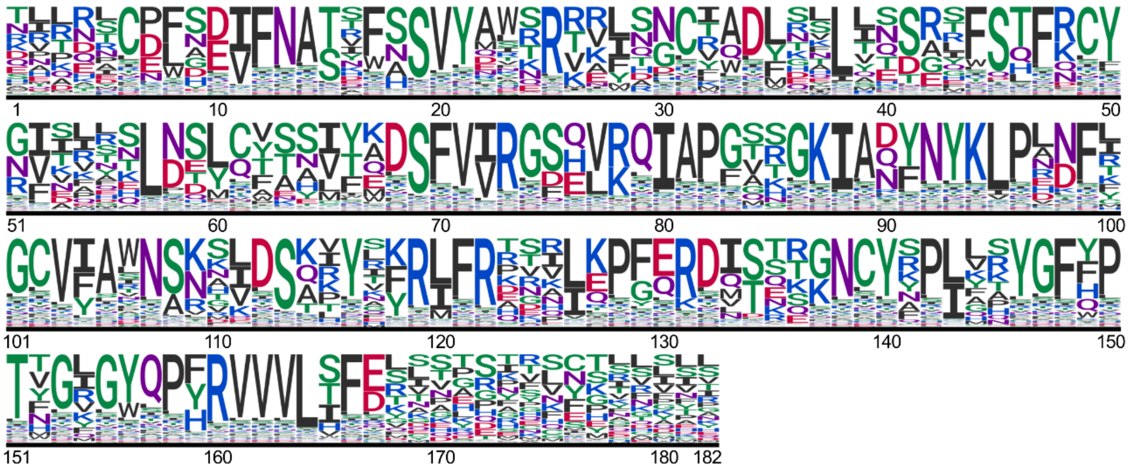

**Supplementary Figure S5. Among-site compositional heterogeneity of the SS-SARS model.** Site-specific estimated equilibrium frequencies estimated by the SS-SARS model are represented from the most frequent to the least amino acid state.

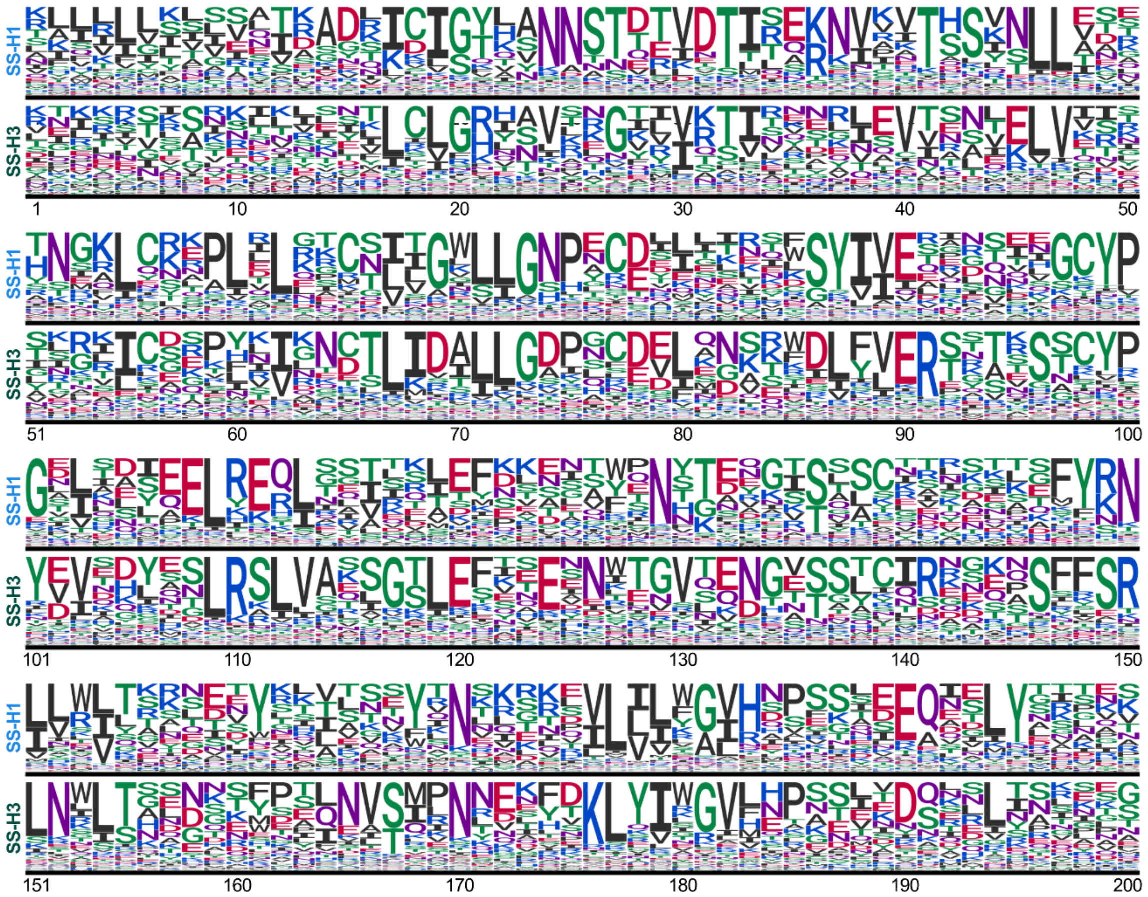


**Supplementary Figure S6. Compositional heterogeneity of the HA-derived SS models.** Site-by-site comparison of the site-specific equilibrium frequencies estimated by the SS-H1 (top) and SS-H3 (bottom) models. For simplicity, only the first 200 out of 550 sites of the protein are shown.

**Supplementary Table S2. Analysis of the association between the disagreements among SS and SH reconstructions and reconstruction ambiguity by maximum parsimony (MP)**

| System/MP reconstruction | Comparison between reconstructions | |
|---|---|---|
| | Disagreement | Identical |
| SR (SS-ER vs JTT) | | |
| Ambiguous | 31 | 130 |
| Unambiguous | 27 | 4980 |
| RBD (SS-SARS vs WAG) | | |
| Ambiguous | 58 | 70 |
| Unambiguous | 13 | 6229 |
| HA (SS-H1 vs FLU) | | |
| Ambiguous | 461 | 1347 |
| Unambiguous | 401 | 47291 |

| System | Odds of disagreement | | Odds ratio‡ | p-value |
|---|---|---|---|---|
| | if MP-ambiguous | if MP-unambiguous | | |
| SR* | 0.238 | 0.005 | 43.98 (25.51-75.82) | $< 2.2 \times 10^{-16}$ |
| RBD* | 0.829 | 0.002 | 397.01 (208.10-757.40) | $< 2.2 \times 10^{-16}$ |
| HA† | 0.342 | 0.008 | 40.36 (34.93 - 46.63) | $< 2.2 \times 10^{-16}$ ($\chi^2$ = 6175, df = 1) |

*Top*, 2x2 contingency tables of classified reconstructions according to their identity between substitution models and their reconstruction ambiguity by parsimony (unambiguous if only one MP state, ambiguous if more than one MP state). *Bottom*, summary table of the strength of association analysis between the presence of disagreements among models and ambiguity by parsimony. Statistically significant p-values that approximate zero are commonly reported as $< 2.2 \times 10^{-16}$ by R packages.

‡ Odds ratio 95% confidence interval in parenthesis

* Significance was assessed using the Fisher's exact test due to the small number of reconstructions with disagreements between models

† Significance was measured using the two-sided Pearson's chi-square test due to its large sample size, $\chi^2$ statistic and degrees of freedom (df) in parenthesis

**Supplementary Table S3. Contingency tables and statistical analysis of the association between ambiguous reconstructions obtained by maximum likelihood (ML) and ambiguous reconstructions obtained by maximum parsimony (MP)**

| MP reconstruction | ML reconstruction | |
|---|---|---|
| | Ambiguous | Unambiguous |
| SR (SS-ER vs JTT) | | |
| Ambiguous | 30 | 98 |
| Unambiguous | 13 | 5027 |
| RBD (SS-SARS vs WAG) | | |
| Ambiguous | 59 | 69 |
| Unambiguous | 7 | 6235 |
| HA (SS-H1 vs FLU) | | |
| Ambiguous | 892 | 848 |
| Unambiguous | 755 | 47005 |

| System | Odds of ML-ambiguous | | Odds ratio | p-value |
|---|---|---|---|---|
| | if MP-ambiguous | if MP-unambiguous | | |
| SR* | 0.306 | 0.003 | 118.37 (59.92 - 233.85) | $< 2.2 \times 10^{-16}$ |
| RBD* | 0.855 | 0.001 | 761.62 (335.90 – 1726.93) | $< 2.2 \times 10^{-16}$ |
| HA† | 1.052 | 0.016 | 65.49 (58.18 – 73.72) | $< 2.2 \times 10^{-16}$ ($\chi^2$ = 12868, df = 1) |

Top, 2x2 contingency tables of classified reconstructions according to their reconstruction ambiguity by maximum likelihood (ML) and their reconstruction ambiguity by maximum parsimony (MP). Bottom, summary table of the strength of association analysis between ambiguously reconstructed states by ML and ambiguity by parsimony.

‡ Odds ratio 95% confidence interval in parenthesis

* Significance was assessed using the Fisher's exact test due to the small number of reconstructions with disagreements between models

† Significance was measured using the two-sided Pearson's chi-square test due to its large sample size, $\chi^2$ statistic and degrees of freedom (df) in parenthesis

**Supplementary Table S4. Analysis of the sites' rate of evolution and sequence entropy and their relationship with the similarity between SS and SH reconstructions**

| System | Sites with identical reconstructions | Sites with reconstruction disagreements | Parameter | D | p-value |
|---|---|---|---|---|---|
| SR (SS-ER vs JTT) | 5110 | 58 | Rate<br>Entropy | 0.4878<br>0.5208 | $2.8 \times 10^{-12}$<br>$6.1 \times 10^{-14}$ |
| RBD (SS-SARS vs WAG) | 6299 | 71 | Rate<br>Entropy | 0.7296<br>0.6222 | $< 2.2 \times 10^{-16}$<br>$< 2.2 \times 10^{-16}$ |
| HA (SS-H1 vs FLU) | 48638 | 862 | Rate<br>Entropy | 0.4249<br>0.4083 | $< 2.2 \times 10^{-16}$<br>$< 2.2 \times 10^{-16}$ |

The relationship between the similarity of SS and SH reconstructions and the strength of phylogenetic signal was evaluated by comparing the frequency distributions of reconstructions when classified by their site's estimated rate of evolution or state diversity (expressed in Shannon entropy values given the alignment data). A two-sample Kolmogorov-Smirnov (KS) test was used to determine if sites with disagreements have the same distribution as sites with identical reconstructions given their rate of evolution and sequence entropy (null hypotheses). The KS test sample size (identical + disagreeing sites), statistic (D) and p-values are reported.

**Supplementary Table S5. Reconstruction accuracy percentage to the true ancestral sequence in the simulated datasets**

| System | Model | Simulated branch length (substitutions/site) | | | | | |
|---|---|---|---|---|---|---|---|
| | | 0.05 | 0.1 | 0.2 | 0.4 | 0.8 | 1.6 |
| SR (n = 2000) | SS-ER† | 98.94 ± 1.19 | 96.8 ± 2.07 | 91.55 ± 3.28 | 82.31 ± 4.51 | 70.69 ± 5.29 | 58.54 ± 5.64 |
| | SS-MR | 98.98 ± 1.16 | 96.87 ± 2.01 | 91.8 ± 3.16 | 82.92 ± 4.28 | 71.84 ± 4.98 | 60.17 ± 5.06 |
| | JTT | 98.88 ± 1.19 | 96.69 ± 2.06 | 91.44 ± 3.33 | 82.39 ± 4.32 | 71.4 ± 4.89 | 60.16 ± 5.1 |
| | Poisson | 98.66 ± 1.29 | 96.06 ± 2.22 | 90.43 ± 3.32 | 81.48 ± 4.35 | 70.59 ± 4.86 | 59.32 ± 5.03 |
| RBD (n = 1000) | SS-SARS† | 98.85 ± 0.8 | 96.73 ± 1.31 | 92.03 ± 2 | 85.09 ± 2.76 | 76.85 ± 3.17 | 68.29 ± 3.51 |
| | WAG | 98.72 ± 0.83 | 96.52 ± 1.37 | 91.73 ± 2.06 | 84.83 ± 2.74 | 76.56 ± 3.12 | 67.84 ± 3.37 |
| | Poisson | 98.37 ± 0.92 | 95.81 ± 1.47 | 90.83 ± 2.15 | 83.86 ± 2.79 | 75.69 ± 3.1 | 67.23 ± 3.4 |
| HA (n = 200) | SS-H1† | 99.31 ± 0.37 | 97.46 ± 0.68 | 92.88 ± 1.2 | 83.94 ± 1.55 | 71.46 ± 1.88 | 57.84 ± 2.03 |
| | SS-H3 | 99.28 ± 0.38 | 97.37 ± 0.72 | 92.76 ± 1.22 | 83.83 ± 1.58 | 71.05 ± 1.91 | 56.91 ± 1.93 |
| | FLU | 99.24 ± 0.39 | 97.28 ± 0.73 | 92.53 ± 1.26 | 83.59 ± 1.59 | 70.77 ± 2 | 56.4 ± 2.01 |
| | Poisson | 98.9 ± 0.45 | 96.39 ± 0.82 | 90.76 ± 1.3 | 81.44 ± 1.69 | 68.4 ± 1.91 | 54.9 ± 1.94 |

Average and standard deviation of the accuracy of the reconstructions for each model. The number of reconstructed sequences per simulated branch length in shown in parenthesis.
† Model used to simulate for the protein family system

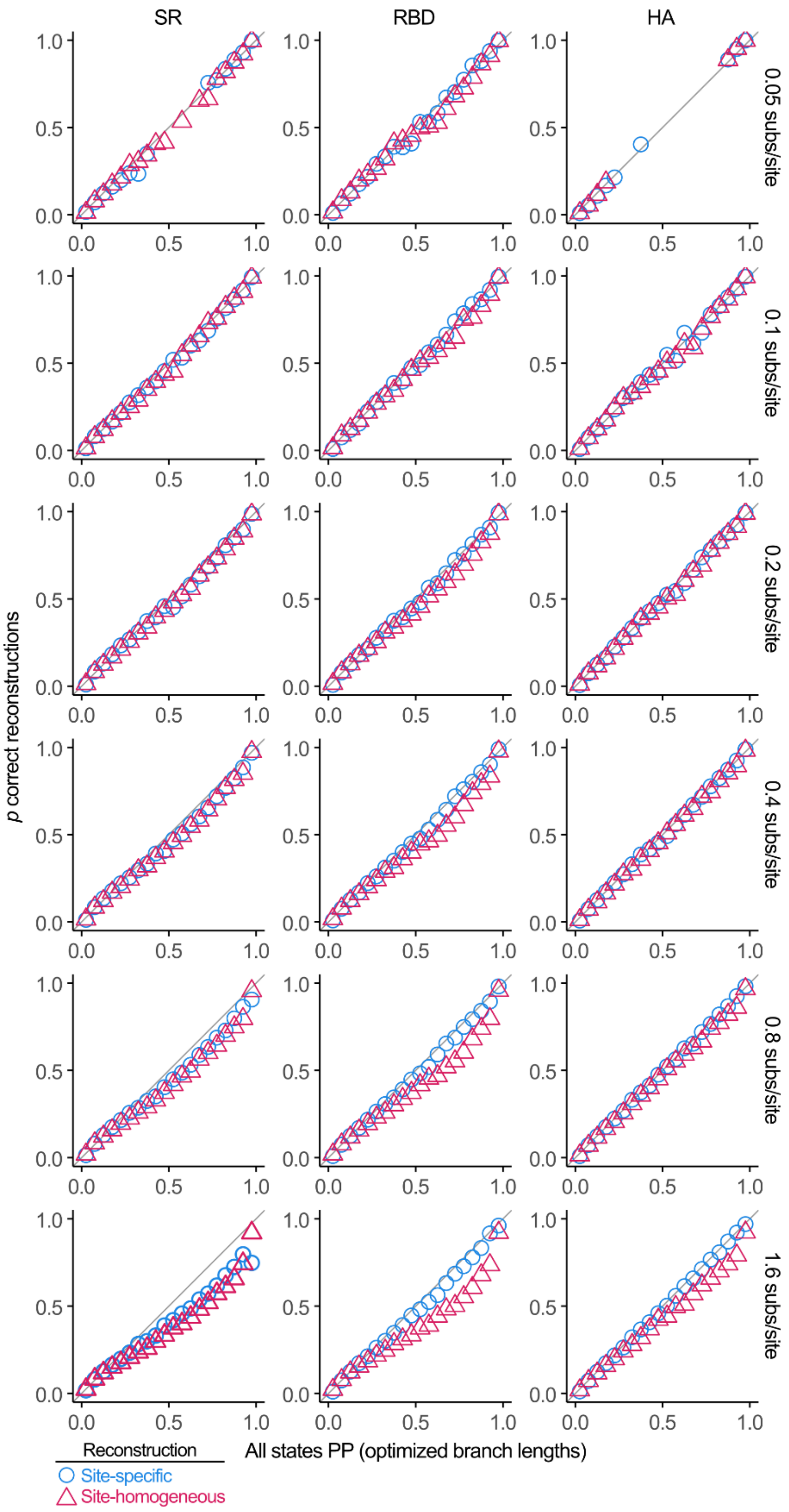


**Supplementary Figure S7. Posterior probability estimation at all simulated branch length conditions.** All possible ancestral states were binned by PP in 5% increments. Only bins with more than 200 elements are shown. Each shape plots the fraction of correct states among reconstructions in a PP bin.

**Supplementary Table S6. Analysis of the association between the disagreements among SS reconstructions and reconstruction ambiguity by maximum parsimony (MP)**

| MP reconstruction | Comparison between reconstruction | |
|---|---|---|
| | Disagreement | Identical |
| SR (SS-ER vs SS-MR) | | |
| Ambiguous | 23 | 138 |
| Unambiguous | 17 | 4990 |
| HA (SS-H1 vs SS-H3) | | |
| Ambiguous | 489 | 1319 |
| Unambiguous | 421 | 47271 |

| System | Odds of disagreement | | Odds ratio | p-value |
|---|---|---|---|---|
| | if MP-ambiguous | if MP-unambiguous | | |
| SR* | 0.167 | 0.003 | 48.92 (25.56 – 93.65) | $< 2.2 \times 10^{-16}$ |
| HA† | 0.371 | 0.009 | 41.63 (36.14 – 47.95) | $< 2.2 \times 10^{-16}$ ($\chi^2 = 6593$, df = 1) |

Top, 2x2 contingency tables of classified reconstructions according to their identity between substitution models and their reconstruction ambiguity by parsimony. Bottom, summary table of the strength of association analysis between the presence of disagreements among models and ambiguity by parsimony.

‡ Odds ratio 95% confidence interval in parenthesis

* Significance was assessed using the Fisher's exact test due to the small number of reconstructions with disagreements between models

† Significance was measured using the two-sided Pearson's chi-square test due to its large sample size, $\chi^2$ statistic and degrees of freedom (df) in parenthesis

**Supplementary Table S7. Analysis of the sites' rate of evolution and sequence entropy and their relationship with the similarity between SS reconstructions**

| System | Sites with identical reconstructions | Sites with disagreements | Parameter | D | p-value |
|---|---|---|---|---|---|
| SR (SS-ER vs SS-MR) | 5128 | 40 | Rate | 0.4562 | $1.34 \times 10^{-7}$ |
| | | | Entropy | 0.4734 | $3.76 \times 10^{-8}$ |
| HA (SS-H1 vs SS-H3) | 48590 | 910 | Rate | 0.3902 | $< 2.2 \times 10^{-16}$ |
| | | | Entropy | 0.3721 | $< 2.2 \times 10^{-16}$ |

The relationship between the similarity of reconstructions using different SS models and the strength of phylogenetic signal was evaluated by comparing the frequency distributions of reconstructions when classified by their site's estimated rate of evolution or sequence entropy. A two-sample KS test was used to determine if sites with disagreements have the same distribution as sites with identical reconstructions given their rate of evolution and sequence entropy (null hypotheses). The KS test sample size (identical + disagreeing sites), statistic (D) and p-values are reported.
